# High-throughput ML-guided design of diverse single-domain antibodies against SARS-CoV-2

**DOI:** 10.1101/2023.12.01.569227

**Authors:** Christof Angermueller, Zelda Mariet, Ben Jester, Emily Engelhart, Ryan Emerson, Babak Alipanahi, Charles Lin, Colleen Shikany, Daniel Guion, Joel Nelson, Mary Kelley, Margot McMurray, Parker Shaffer, Cameron Cordray, Samer Halabiya, Zachary Mccaw, Sarah Struyvenberg, Kanchan Aggarwal, Stacey Ertel, Anissa Martinez, Snehal Ozarkar, Kevin Hager, Mike Frumkin, Jim Roberts, Randolph Lopez, David Younger, Lucy J. Colwell

## Abstract

Treating rapidly evolving pathogenic diseases such as COVID-19 requires a therapeutic approach that accommodates the emergence of viral variants over time. Our machine learning (ML)-guided sequence design platform combines high-throughput experiments with ML to generate highly diverse single-domain antibodies (VHHs) that bind and neutralize SARS-CoV-1 and SARS-CoV-2. Crucially, the model, trained using binding data against early SARS-CoV variants, accurately captures the relationship between VHH sequence and binding activity across a broad swathe of sequence space. We discover ML-designed VHHs that exhibit considerable cross-reactivity and successfully neutralize targets not seen during training, including the Delta and Omicron BA.1 variants of SARS-CoV-2. Our ML-designed VHHs include thousands of variants 4-15 mutations from the parent sequence with significantly improved activity, demonstrating that ML-guided sequence design can successfully navigate vast regions of sequence space to unlock and future-proof potential therapeutics against rapidly evolving pathogens.

## Introduction

The recent COVID-19 pandemic has highlighted the need for therapeutic strategies that combat fast-evolving pathogens. The swift initial response to the pandemic resulted in multiple therapeutic antibody approvals. However, the SARS-CoV-2 virus evolved rapidly, rendering several approved therapies ineffective against today’s predominant variants^1^. Additionally, developing new antibody therapeutics remains costly, time-consuming, and plagued with risk: even antibodies with the right binding profile may ultimately fail due to developability or immunogenicity issues^2–6^. One potential solution is a platform that generates large sets of diverse antibody candidates with the required binding activity, under the hypothesis that diversity provides a buffer against downstream functional requirements that cannot be measured at scale, such as developability, immunogenicity, and neutralization of new viral variants. Moreover, the rapid growth of multispecific therapeutics provides the opportunity to combine the advantages of diverse candidates^7^, however, designing large sets of diverse antibodies remains an unsolved challenge^3,8^.

High-throughput platforms, such as phage and yeast display, can screen libraries containing 10^9^-10^11^ antibody variants^9,10^. But even with a library size of 10^11^, restricting mutations to the ∼30 positions in the complementarity determining regions (CDRs) of a VHH, we can only exhaustively explore up to ∼5 mutations from a parent sequence. Achieving sufficiently diverse candidate VHHs to hedge against new viral variants and other downstream requirements would be greatly accelerated by the ability to search beyond this radius. Moreover, a limitation of existing platforms is that they typically screen one property at a time, such as binding to a single target, making it difficult to optimize multiple properties in parallel at scale, such as cross-reactive binding to multiple viral variants.

Machine learning (ML)-guided antibody design combined with a high-throughput experimental platform has the potential to address these limitations^3,8^. In this approach, a model is trained to capture the relationship between amino acid sequences and multiple experimental measurements, such as binding affinity to multiple targets (Figure 1). This model can predict the effect of novel mutations *in silico* and prioritize new antibody sequences for experimental validation. Iterative design-build-test cycles improve the model and extend the range of experimental properties that it captures. While existing studies of ML-guided antibody design have shown promising results, most focus on exploring the sequence space up to 3-4 steps from a parent sequence, while multi-round ML-guided optimization campaigns have not been widely explored^11–14^. Additionally, due to a lack of suitable datasets, only a handful of studies^13^ train models using experimental data for more than one functional property.

**Fig. 1.**
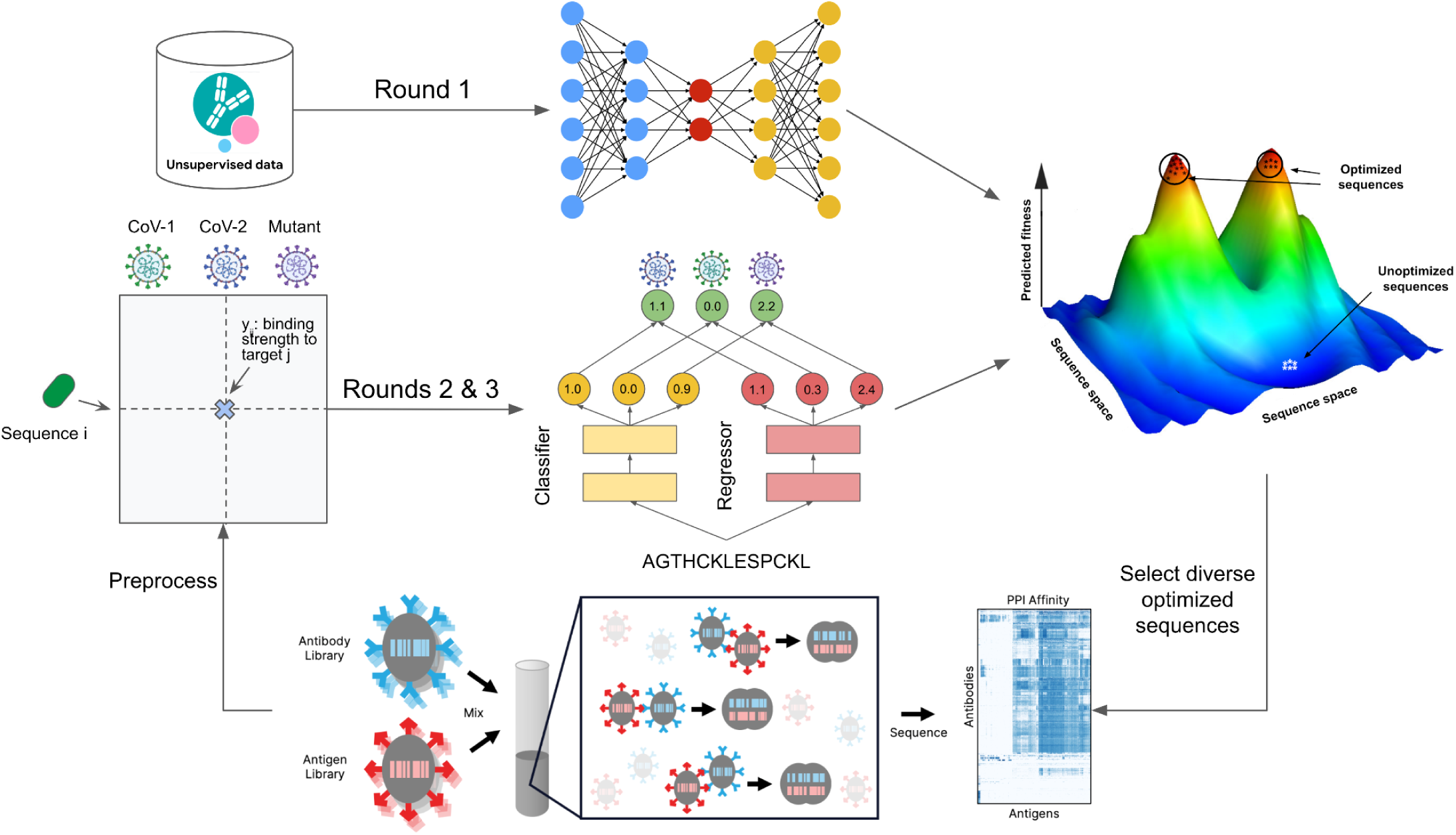
Machine-learning guided antibody optimization. In Round 1, a large set of natural VHH sequences from the OAS database was used to train variational autoencoders (VAEs) to describe the space of viable VHH sequences. The binding of a large set of VHH sequences selected by these models to different coronavirus variants is experimentally characterized using the AlphaSeq assay, and used as training data in round 2 for initial regression and classification models that predict binding strengths given an input VHH sequence. Whereas the classifier predicts whether an input VHH sequence binds to a specific antigen, the regressor predicts continuous binding scores, and the final model output is the product of the classifier and regressor score. The trained model is optimized *in silico* by searching for sequences with high predicted affinities. A diverse subset of optimized sequences is selected and validated experimentally using the AlphaSeq assay. The resulting experimental measurements are added to the training data for round 3 models.

In this work, we use ML to optimize the binding affinity and cross-reactivity of a parent sequence, VHH-72, to a set of coronavirus variants. VHH-72, the precursor to the biologic XVR011, exhibits high affinity binding against SARS-CoV-1 and weaker binding against SARS-CoV-2^15,16^. VHH-72 binds to a conserved epitope of the spike protein receptor binding domain (RBD), a mechanism that enables significant cross-reactivity and reduces susceptibility to escape mutations. To improve cross-reactivity, we carried out three rounds of ML-guided VHH design, combining supervised and unsupervised ML models. We used the AlphaSeq assay^17,18^ to simultaneously measure antibody binding affinity against 23 RBD targets, including SARS-CoV-1 and SARS-CoV-2, eight SARS-CoV-2 single point mutants, two SARS-CoV-2 triple mutants, and eleven distantly related sarbecoviruses. The assay also includes six positive and three negative control targets(Methods). As a control, we ran a parallel “baseline” campaign using a simple additive model to combine mutations from the best variants found so far.

We found that ML-guided antibody design identifies highly cross-reactive sequences with significantly improved binding activity against SARS-CoV-2 with a success rate ∼70%, compared to <23% for the baseline. Our best sequences improve SARS-CoV-2 RBD binding affinity and pseudovirus neutralization by more than 50-fold. Strikingly, these top sequences also improve binding and neutralization against the B.1.617.2 (Delta) RBD variant by 48-fold and 94-fold respectively, although Delta RBD mutations were not seen during training. Moreover, in contrast to VHH-72, 5/19 ML-designed sequences selected for additional testing tightly bind and neutralize the BA.1 (Omicron) RBD, which contains 15 RBD mutations^19^, of which only three (S477N in round 1, K417N and N501Y in round 2) were seen during training. Overall, our results demonstrate that ML-guided optimization for cross-reactivity against multiple targets can yield a plethora of highly diverse sequences with significantly improved binding activity, providing a therapeutic hedge against future pathogen evolution.

## Results

In pursuit of a diverse panel of highly cross-reactive antibodies, we first characterized the binding activity of sequences up to four mutations away from the parental VHH-72, against 21 diverse sarbecovirus RBDs, including SARS-CoV-1, SARS-CoV-2, eight single mutant variants of SARS-CoV-2 and eleven distantly related sarbecoviruses (Extended Data Table 1)^20^. We focused on the 34 CDR positions, in addition to 14 framework region (FWR) positions that were most variable across 542,643 unique VHH sequences from the Observed Antibody Space (OAS) database^21^. To generate training data, we synthesized 12,000 VHH sequences with up to 4 mutations from VHH-72. We include all 864 single mutants (excluding histidine) in the first library, and consider them part of the baseline campaign. To capture dependencies between sequence positions, we used the OAS data to train variational autoencoders (VAEs)^22,23^ and performed *in silico* screening of all double-mutants and 20 million triple and quadruple mutants with mutations confined to CDR1+CDR2 or CDR3 (Methods). We selected 2,000 sequences with the highest VAE likelihood at each distance for synthesis, in addition to 5,136 multi-mutant sequences selected randomly (above a threshold), to increase diversity. The binding activity of each VHH in the library was measured in triplicate using the AlphaSeq assay. To measure assay variability across replicates and quantify statistical significance, 46 separately synthesized copies of the VHH-72 sequence were also included.

Sequences with mutations in CDRs 1/2 typically retained binding activity, while variants with mutations in CDR3 largely lost binding activity against both SARS-CoV-1 and SARS-CoV-2, highlighting the importance of this region (Extended Data Fig. 1). The binding profiles of six single mutants of SARS-CoV-2 correlated with those of the SARS-CoV-2 and SARS-CoV-1 wildtypes (WTs, Extended Data Fig. 1); data from these eight targets were used to train several models that capture the relationship between VHH sequence and binding activity. To take advantage of the large portion of sequences with no measurable binding, we first trained multi-task classification models to predict whether a VHH sequence will bind these targets, and then also trained regression models to predict quantitative binding affinity. The product of the classification and regression predictions for each variant predicts the binding affinity (<u>Methods</u>).

Through cross validation on the first round data, we found that ensembles of light gradient-boosted (LGB) models outperformed ensembles of convolutional neural networks (CNNs) and a baseline linear model (Figure 2A). To design sequences that bind to multiple SARS-CoV variants and are robust to escape mutations, we use the weighted average of LGB-predicted affinities against the eight targets used for model training to score each sequence, and constrain the optimization to a trust region by adding a linear penalty to sequences with more than eight mutations. An ensemble of discrete optimization algorithms, previously shown to find diverse high-scoring sequences in silico^24^, was used to optimize the resulting scoring function. To design baseline sequences, we greedily combined the best mutations or combinations of mutations found in round 1 by (i) combining the mutations found in pairs of top sequences and (ii) adding the best single mutants to each top sequence (Methods).

**Fig. 2.**
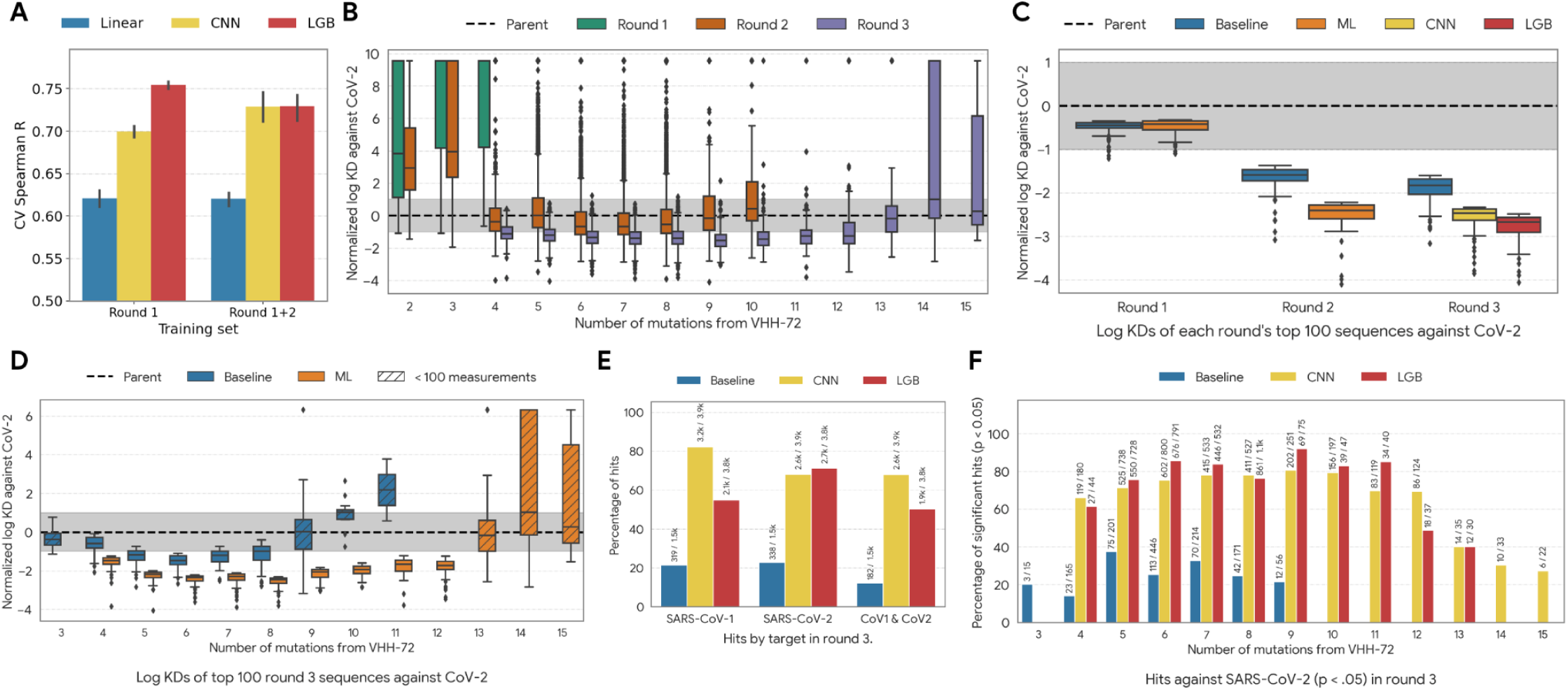
Performance of VHH design methods. (A) 5-fold cross-validation performance quantified by Spearman’s *R* of models trained on data from round 1, or round 1 and 2, averaged across targets. Strong LGB performance motivated the use of this model to design round 2, while the LGB and CNN models were both used to design round 3. (B) Comparison of normalized binding affinities (lower is better) stratified by distance from VHH-72 for sequences from the three designed libraries (note that for clarity, sequences replicated from earlier rounds for calibration are not included in the results reported for later rounds). Grey shading indicates the range of AlphaSeq measurements across wildtype (WT) replicates. (C) AlphaSeq measurements for the best 100 ML-designed and baseline sequences from each round. For round 1 the baseline consists of all single mutant sequences. (D) AlphaSeq measurements for the best 100 ML-designed and baseline sequences at each number of mutations relative to VHH-72. Hashed shading indicates that <100 sequences were measured, in which case all measurements were included. (E) Fraction of ML-designed and baseline sequences from round 3 that significantly improve binding compared to VHH-72 (p < 0.05, Mann-Whitney U-test, Bonferroni correction) while maintaining WT SARS-CoV-1 binding activity. (F) Fraction of round 3 sequence designs with significantly improved binding against SARS-CoV-2 compared to VHH-72 (*p* < 0.05, Mann-Whitney *U*-test, Bonferroni correction) as a function of number of mutations from VHH-72; ML-designed sequences with up to 15 mutations had significantly improved binding activity, as did ∼80% (3877/4853) of ML-designed sequences with 6-10 mutations.

To experimentally evaluate the second library, we retained 3 targets (SARS-CoV-1, SARS-CoV-2, and SARS-CoV-2_R408I) and added three newly emerged SARS-CoV-2 variants of concern (B.1.1.7 [Alpha], P.1 [Gamma], and B.1.351 [Beta], Extended Data Table 1). Across these targets, two substitutions from B.1.351 (K417N and N501Y) appear in the Omicron BA.1 RBD and neither of the Delta RBD mutations are present. Reducing the number of targets to six allowed us to increase the number of VHH variants tested without increasing the number of target-VHH interactions, thus maintaining the resolution of the AlphaSeq assay. We synthesized 28,260 ML-designed VHH sequences, 5,821 baseline sequences, 51 copies of VHH-72, and 150 VHHs from round 1 with stratified binding affinities in triplicate for calibration. Experimentally, 25.8% of ML-designed sequences improve binding to SARS-CoV-2 by at least 1 IQR, compared to 5.4% of baseline sequences. In contrast to round 1, ML-designed sequences with CDR3 mutations were more likely to improve binding than those with mutations in CDR1 and CDR2 only (Extended Data Fig. 2).

**Table 1.** Number of viable *k*-mutants of the parent VHH-72, before and after selection of mutable positions.

|  |  |  |  |  |
| --- | --- | --- | --- | --- |
| # Mutations | 1 | 2 | 3 | 4 |
| # Sequences | 2.4k | 2.8M | 2.2B | 1.3T |
| # Sequences after position filtering | 864 | 88k | 8M | 512M |

To design the final library, we again combined classification and regression models, trained on data from the two previous rounds. CNN and LGB ensembles achieved similar cross-validation performance (Figure 2A, Extended Data Fig. 3), so both were used to design the third library, with the goal of increasing the diversity of the designed sequences. To further increase diversity, we clustered candidate sequences and kept the top 4000 scoring sequences from each cluster and model. We designed 1500 baseline sequences by recombining the mutations from the best baseline sequences found so far. To calibrate measurements with previous rounds, we included 696 VHHs with stratified binding affinities from round 1 and 49 copies of VHH-72.

To ensure quantitative resolution for tight binders we limited targets to SARS-CoV-1 and SARS-CoV-2 WT (Extended Data Table 3), replicated each sequence three times, and increased the number of technical AlphaSeq replicates to six. Both ML-designed and baseline sequences bound more tightly compared to previous rounds (Figure 2B-C, Extended Data Fig. 4). Overall 5,361/7,716 (69.5%) ML-designed sequences with up to 15 substitutions showing significant (*p* <= 0.05 Mann-Whitney *U*-test, Bonferroni correction) improvement over VHH-72 against SARS-CoV-2 WT, compared to 338/1496 (22.6%) baseline sequences with up to nine substitutions (Figure 2D, E, F). We note that previously reported VHH-72 variants with improved binding affinity that have up to 8 substitutions^15,25,26^. Overall these results provide evidence that the ML models extrapolate accurately to identify functional sequence variants with multiple mutations.

To confirm the results of our high-throughput AlphaSeq measurements, we used biolayer interferometry (BLI, Methods) to measure binding affinities for 21 VHH sequences, including VHH-72 and the best single-mutant sequence (Seq12). To select 19 additional sequences, we first excluded sequences that bound to negative control targets in any replicate or had mutations to cysteine, then built three sets of 200 sequences with (i) the smallest Benjamini-Yekutelli adjusted p-value, (ii) the tightest affinity (median-aggregated over replicates), and (iii) the tightest affinity (mean-aggregated over replicates). All 68 sequences in the intersection of these three sets were ML-designed, reflecting strong ML-guided design performance compared to baseline. Finally, we clustered these 68 sequences into 10 groups and selected the 1-2 sequences with smallest adjusted p-value from each group to yield 19 ML-designed sequences.

For each sequence, we measured protein expression and binding affinity to the SARS-CoV-1 and SARS-CoV-2 RBDs (Methods, Extended Data Fig. 5). In agreement with previous reports, we found that VHH-72 bound >5x weaker to SARS-CoV-2 than to SARS-CoV-1^15,26^. Overall, 13 VHH sequences (Seqs 1-12 and VHH-72), showed high protein expression (>10 mg/L) in *E. coli.* Notably, all 19 ML-designed VHH sequences showed >2-fold improvement in SARS-CoV-2 binding affinity, with the best candidate reporting ∼50-fold improvement (Figure 3A, Extended Data Figs. 5, 6). ML-designed sequences also retained or improved binding to SARS-CoV-1 by up to ∼8-fold. The greater improvement in binding affinity against SARS-CoV-2 likely reflects already strong binding to CoV-1 and the additional weight assigned to SARS-CoV-2 targets during model-based optimization (Methods).

**Fig. 3.**
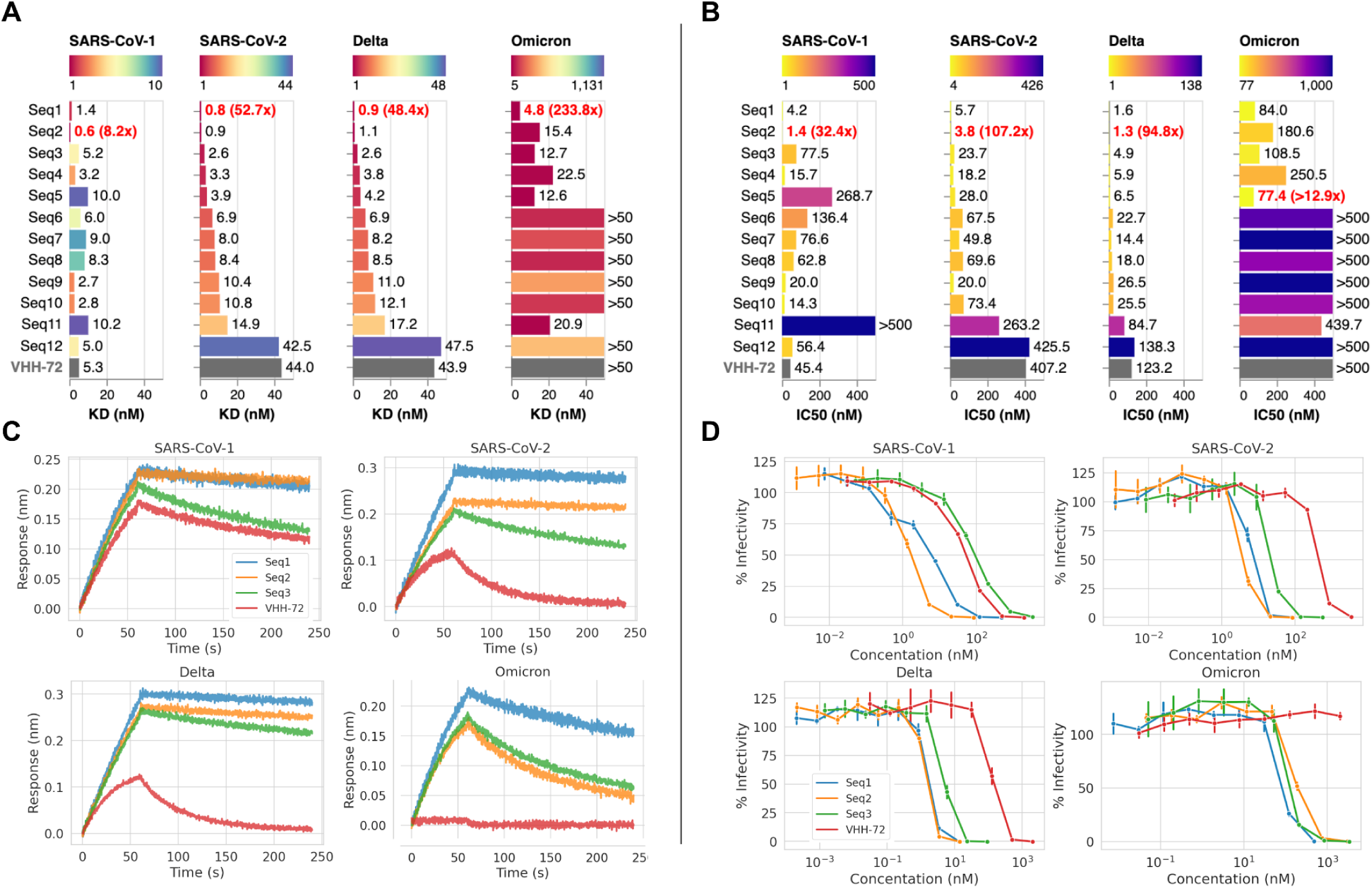
Biolayer interferometry (BLI) and pseudovirus neutralization measurements of selected VHHs against CoV targets. (A) BLI binding strengths in KD (nM) of representative best ML-designed sequences with high expression levels (Seqs 1-11), the best single mutant variant (Seq 12) and VHH-72 for SARS-CoV-1, SARS-CoV-2, Delta, and Omicron. A lower value indicates stronger binding. The binding strength and fold improvement of the best binding sequence is highlighted in red. (B) Pseudovirus neutralization potency (IC50) in KD (nM) corresponding to the sequences and targets shown in A. (C-D) BLI binding curves and neutralization curves of Seq1-3 and VHH-72.

We further tested the 13 high expression VHH sequences in pseudovirus neutralization assays against SARS-CoV-2 RBD, finding that the 11 ML-designed sequences (Seqs 1-11) improved neutralization up to 100-fold (Figure 3B, Extended Data Fig. 5). In contrast, the single mutant baseline sequence (Seq 12) showed no improvement in binding affinity or neutralization. We also tested neutralization of the Delta and Omicron BA.1 variants, finding that the best ML-designed sequences improved neutralization of the Delta variant by ∼100x (Figure 3B). Importantly, while VHH-72 displays micromolar binding to the Omicron BA.1 RBD and has no detectable pseudovirus neutralization, our best ML-designed sequences display nanomolar binding and sub-100 nanomolar neutralization (Figure 3). This is despite the fact that the Omicron BA.1 RBD is 14 mutations away from the closest binding target used to generate training data for ML models. Together, these results show that ML-guided design can discover diverse VHH sequences that improve binding and neutralization activity against emerging pathogenic virus strains not seen in model training.

Figure 4A shows the sequences characterized by BLI that expressed well, while Figure 4B shows the sequence distribution of the 100 ML-designed sequences with tightest binding to SARS-CoV-2 RBD WT as measured using AlphaSeq. We find enrichment for previously reported mutations, including S56A ^16^, S56G, T99V, and V100W^25^, and S56M, L97W^26^. Notably, we also find a high prevalence of novel mutations, including G55R/K in CDR2, A94R, L97Y/W/F, and V100Y in CDR3. All 5/11 tested variants with neutralization activity against Omicron BA.1 contain L97Y/F, four contain G55R/K, three contain A94R and two contain V100Y. Our model predicts that combining favorable single-mutants will increase binding (Extended Data Fig. 7), while most CDR3 mutations are predicted to strongly decrease binding, highlighting the challenge of optimizing this region. We used the model to probe the predicted epistatic effects of pairs of newly discovered mutations (Methods). The model predicts weak positive epistatic effects between L97 and A94, and weak negative epistatic effects between L97 and G55, T99, or V100 (Extended Data Fig. 8). Overall these results provide evidence that the ML models were able to identify and combine novel mutations to increase binding efficacy.

**Fig. 4.**
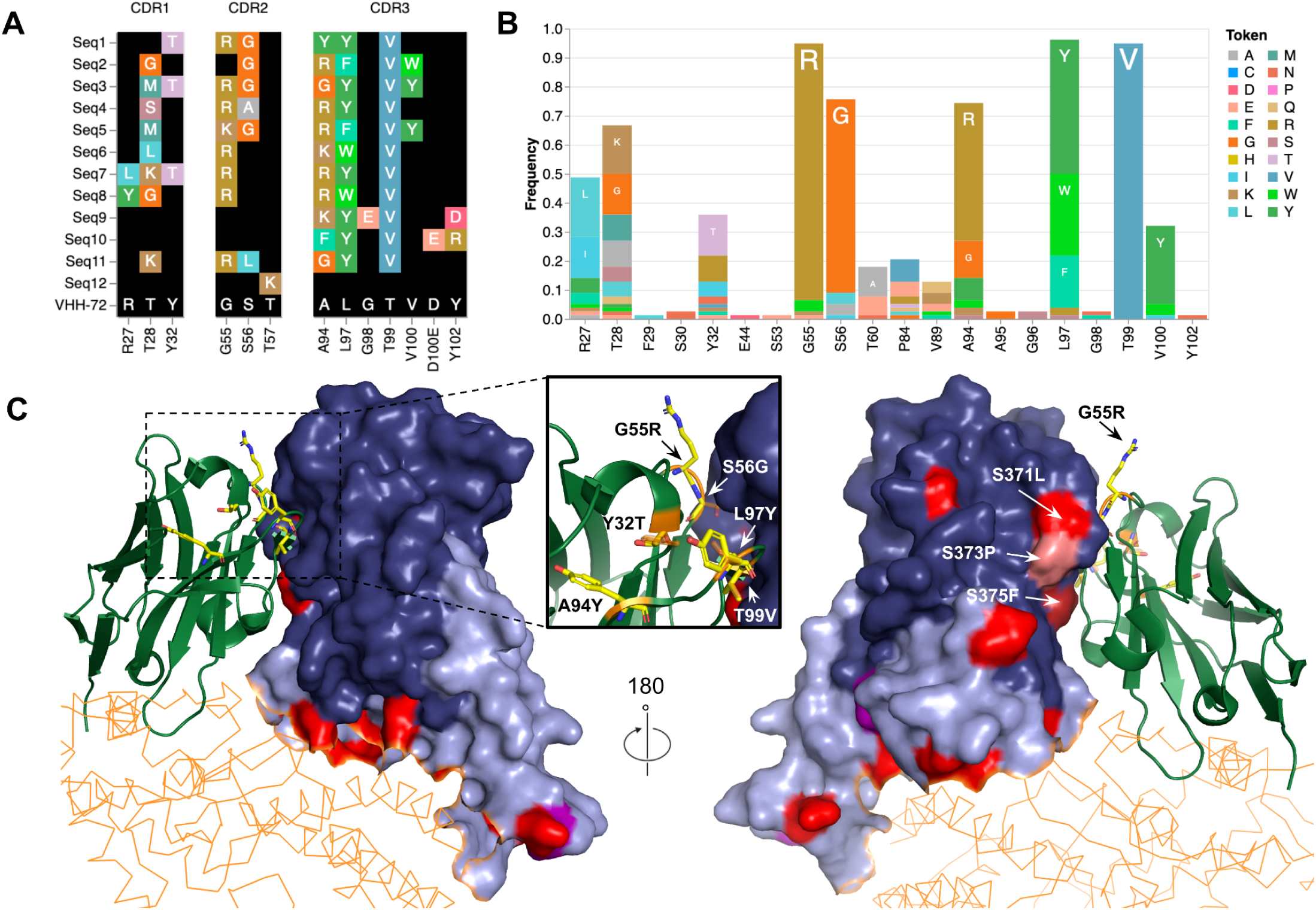
Machine learning identified variants with multiple mutations that improve binding. (A) mutations of the best ML-designed sequences validated by BLI and pseudovirus neutralization relative to VHH-72. (B) Sequence logo of the best 100 ML-designed sequences measured using AlphaSeq. Sequences are enriched for mutations G55R, S56G, A94R, L97Y, L97W, T99V and V100Y. (C) Front- and back view of VHH-72 (green; PDB 6waq) bound to the receptor-binding domain (RBD) of SARS-CoV-2 (blue; PDB 6m17) in conjunction with the Human ACE2 receptor (orange; PDB 6m17). Mutations of the best ML designed sequences Seq1 are modeled and shown as yellow sticks. The receptor-binding motif (RBM) of the RBD is shown in light-blue. Mutations of the Delta and Omicron BA.1 variant are highlighted in purple and red, respectively. The zoom-in view shows the mutations of the best designed sequence *Seq1*, with wildtype residues of VHH-72 highlighted in orange and target residues of *Seq1* in yellow.

To better understand the improved potency of our designed sequences, we probed the structural context. In contrast to most therapeutic antibodies, which bind the receptor-binding motif (RBM) of the RBD, VHH-72 binds to a cryptic epitope of SARS-CoV-2 through a hydrogen-bonding network involving CDR2 and CDR3^15^. This mechanism instills resistance to frequently observed escape mutations in the RBM region, such as L452R, and T478K in the Delta variant. However, five Omicron RBD mutations are located outside of the RBM, including three mutations (S371L, S373P, S375F) at the VHH-72 epitope, none of which were seen during model training. Our top five BLI-tested sequences (Seq1 - Seq5, Figure 3) feature G55R/K, S56G, L97Y/W/F and T99V, which fall within the paratope if the interface between WT VHH-72 and the SARS-CoV-1 RBD is maintained (Figure 4C). All of these mutations are highly prevalent among the 100 tightest binding ML-designed sequences, suggesting they may be involved in increasing binding affinity.

Fig. 5A compares quantitative binding predictions from the LGB model trained on data from rounds 1 and 2 against measurements for round 3 sequences against the SARS-CoV-2 WT target, see Extended Data Fig. 9 for other models and training sets, summarized in Figure 5B. These correlations are lower than the retrospective correlations from cross validation on the training data (Fig. 2A), reflecting the challenging nature of prospective design, although the gap closes as the amount of training data increases. Figure 5C stratifies this data by distance from VHH-72 to show that strong prospective design performance is maintained over distance, indicating that the models are able to accurately extrapolate beyond the training data. Model performance at predicting whether sequences bind more tightly than VHH-72 is summarized in Figure 5D, and stratified by distance from VHH-72 in Extended Data Fig. 10. We find that the models generalize well at this classification task, with the LGB and CNN models consistently outperforming a baseline linear model. Comparison with Extended Data Fig. 3 suggests that there is little difference between prospective and retrospective performance for the ML models trained on rounds 1 and 2, in contrast to the baseline linear model.

**Fig. 5.**
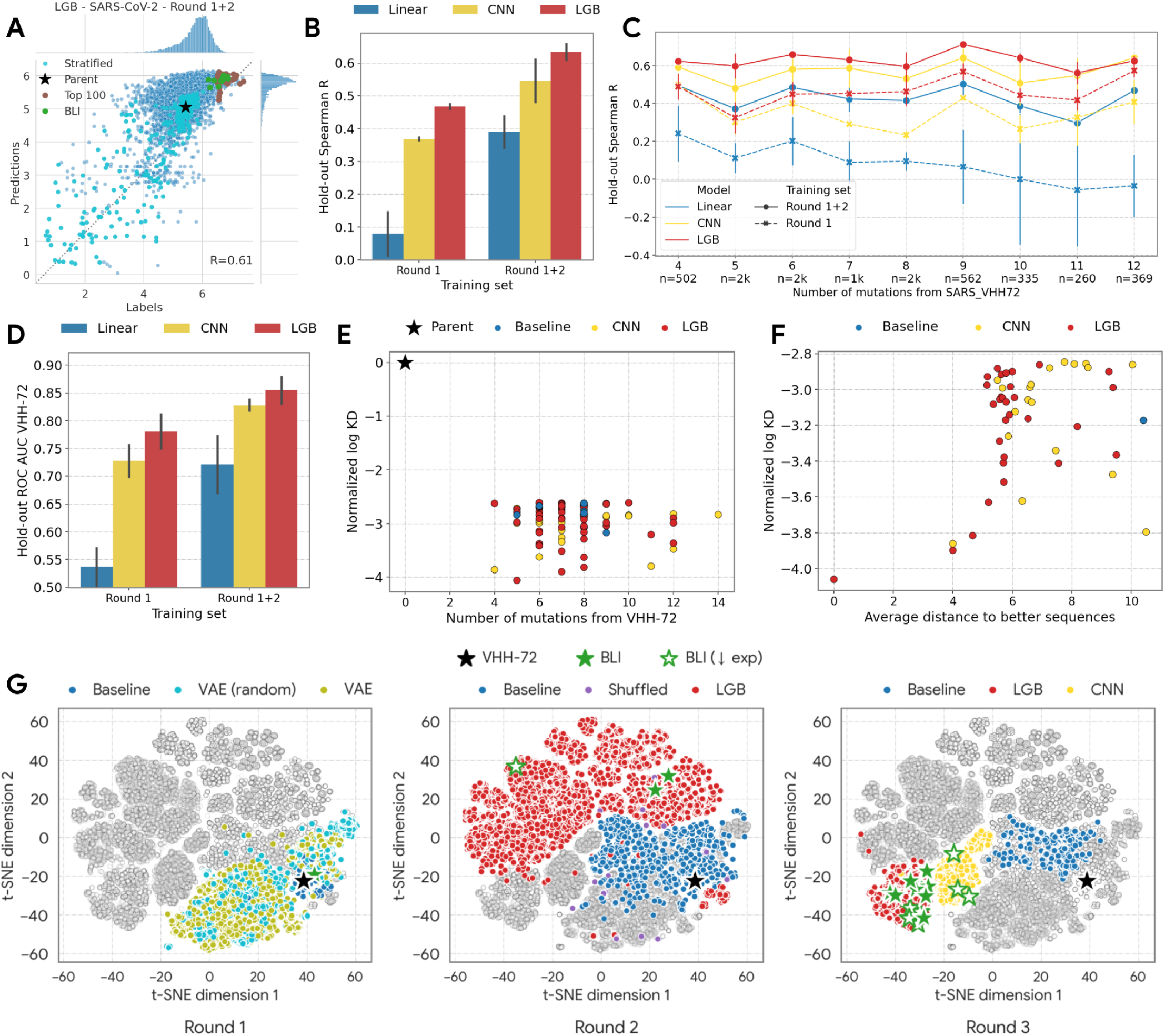
Prospective experimental validation of ML models and diversity of ML-designed sequences: (A) Scatterplot of predicted normalized log KD values vs. experimental measurements for binding of round 3 sequences to SARS-CoV-2. Predictions made using the LGB model trained on round 1 + round 2 data. Each dot corresponds to one sequence assayed in round 3. Measurements and predictions correspond to binding affinities (higher is better) obtained by normalizing AlphaSeq log KD values and computing the difference from the maximum observed measurement. We highlight VHH-72, the set of sequences with stratified log KD values tested in every round, sequences validated by BLI, and the 100 sequences with lowest AlphaSeq log KD values. (B) Prospective correlation (Spearman’s *R*) between model predictions and experimentally measured binding across ML-designed sequences tested in round 3, using models trained on round 1, or round 1 + round 2 data. Error bars show the variation over wild-type CoV-1 and CoV-2. (C) Data from (B) stratified by the distance of ML-designed sequences to VHH-72. (D) ROC AUC score for prospective classification of whether ML-designed sequences tested in round 3 bind more tightly than VHH-72 (see text for details). (E) Normalized affinity (lower is better) of the top 100 round 3 sequences measured using AlphaSeq as a function of the number of mutations from VHH-72. Note the high-affinity ML-designed sequences with 10-14 mutations from VHH-72. (F) The tightest-binding sequences are highly diverse: here we show the average number of mutations between each sequence and sequences with higher affinity for the best 50 sequences discovered in round 3. (G) Two-dimensional t-SNE embedding of all sequences designed over three optimization rounds (sub-panels), highlighted by the corresponding design strategy. Markers highlight VHH-72 and the 19 ML-designed sequences validated by BLI, all of which had >2x improvement in SARS-CoV-2 binding affinity. Those with high (>10 mg/L) expression yield are solid green stars, while low expression yield is open green stars (BLI ↓exp).

The ability to design diverse tight binding sequences is important because it provides a buffer against downstream or future requirements, such as the ability to neutralize newly evolved viral strains. Fig. 5E shows that the 100 tightest binding sequences each have 4-14 mutations from VHH-72. For each of these sequences, the average pairwise distance to sequences that bind more tightly is at least four and can be as high as 10 (Figure 5F), confirming that the models successfully designed highly diverse tight-binding sequences. This diversity is further illustrated by the t-SNE projections of designed sequences over all three rounds in Figure 5G. We note that including CNN- and LGB-designed sequences does increase diversity as anticipated. Overall, our data shows that ML-guided sequence design can leverage models that accurately extrapolate beyond the training data to successfully design diverse sequences with significantly improved binding affinity and the ability to neutralize novel targets.

## Discussion

In this study, we demonstrate that ML-guided sequence design successfully optimizes VHH binding activity against multiple coronavirus strains simultaneously. In addition to substantially improving binding to SARS-CoV-2, ML-guided sequence design generates VHH variants with broad cross-reactivity that extends to emerging strains such as Omicron that were not known when the models were trained. The ability to future-proof potential therapeutics by generating highly diverse candidates that simultaneously bind emerging strains while improving binding activity against existing strains presents a crucial advantage when combating a rapidly evolving pathogen.

To our knowledge, this is the first study in which machine learning has been applied to the generation of highly cross-reactive antibody sequences by explicitly combining data against multiple targets at a large scale. Notably, this approach enables multi-target antibody optimization, yielding sequences with increased cross-reactivity and extensive coverage of a pathogen’s mutational landscape. Out of eleven representative sequences that expressed well, five (45%) acquired neutralization activity against Omicron, demonstrating that our protocol is adept at producing therapeutic candidates with heightened resistance to novel target mutations. This approach can be applied to antibody candidate generation for any infectious disease where target diversity is a challenge, resulting in a cocktail of optimized candidates with broad cross-reactivity.

Extended Data Fig. 6 shows significant variation in expression titers even among the handful of sequences subjected to high-resolution screening in this study, suggesting that there may also be substantial variation in developability properties such as stability and solubility. This finding underscores the utility of an optimization strategy that identifies highly diverse sequences with the desired binding cross-reactivity profile, thereby improving the likelihood of isolating a sequence with ideal properties during subsequent bio-developability assessments. Going forwards, models that predict developability properties^27–29^ can be incorporated into the sequence optimization objective. Taken together, the combination of highly diverse functional VHHs with required binding activity and successful multi-objective optimization point to the potential for ML-guided sequence design to significantly lower attrition in the drug discovery process.

A key limitation to the high-throughput cross-reactivity measurement protocol used here is the requirement for target variants to be displayed on the yeast surface, limiting application to proteins that can be expressed using this method and, e.g., do not require humanized post-translational modifications^30^. In such cases additional data collected using alternative high- or low-throughput methods can be incorporated into model training. Additionally, multiple design-build-test cycles present a challenge in scenarios where time is of the essence, as for an emerging pandemic. However, as more data characterizing VHH binding becomes available, modeling advances^31–33^ could reduce this requirement. We hope that others will take advantage of the dataset of 464,541 quantitative antibody-antigen interactions released with this study to address these research challenges.

## Methods

### Design of the 1st library

We constructed the first library of 12,000 VHHs with the goal of exploring the landscape close to the parent sequence VHH-72 to generate training data. For this purpose, we only considered sequences with up to four mutations from VHH-72. To reduce the search space, we restricted mutations to all 34 positions within the 3 CDR regions (8 positions in CDR1, 8 positions in CDR2, 18 positions in CDR3). We also allowed mutations at the 14 FWR positions (2 in FWR1, 5 in FWR2 and 7 in FWR3) that varied the most across the natural VHH sequences used to train VAE models (described below). Synthesis constraints using 200nt oligos restricted multi-mutant variants to contain mutations in either CDR1 and CDR2 (CDR1+CDR2), or CDR3 (FWR positions were only present in single-mutant variants). These constraints mean that variants with mutations in both CDR1 and CDR3, or CDR2 and CDR3 were not allowed in the first round; further narrowing the search space. Lastly, we disallowed mutations to histidine since it is depleted in naturally occurring VHH sequences.

We included all 864 remaining single-mutants, and used an unsupervised approach to select multi-mutants with up to four mutations. Specifically, we trained variational autoencoders (VAEs)^22,23^ on 542,643 natural VHH sequences^34^ from the Observed Antibody Space (OAS) database^21^, which we aligned using the tool ANARCI^35^. We then selected sequences with the highest likelihood under these models. This unsupervised approach does not require VHH-CoV binding measurements. Our VAE model used a fully-connected encoder and decoder architecture with one hidden layer. To increase the diversity of selected sequences and robustness, we trained two VAE models with different hidden (256 vs. 128) and latent (128 vs. 64) layer sizes.

We enumerated mutants with 2-4 mutations and selected the 6,000 sequences with the maximum likelihood under the two trained VAE models. Sequences that bind to CoV2 may be underrepresented in the set of natural OAS sequences that we used for training VAE models and therefore not be amongst the sequences with the maximum likelihood. To address this, we also selected 5,136 sequences using reservoir sampling. Specifically, we selected a sequence if its VAE score s was above a predetermined threshold T=-95, or uniformly with probability N(s) / N(T), where N is the normal distribution with mean and standard deviation of the VAE scores.

The top-scoring sequences had mutations in CDR3 ∼3.5x more often than in CDR1+2. We further ensured that the number of sequences with mutations in CDR1 was equal to the number of sequences with mutations in CDR2, and split the budget uniformly between variants with 2, 3, and 4-mutations. Altogether, we included

- 558 sequences with mutations in CDR1 only
- 554 sequences with mutations in CDR2 only
- 2,189 sequences with mutations in CDR1 and CDR2
- 7,835 sequences with mutations in CDR3 only

### Data preprocessing

Within a given AlphaSeq experiment, each VHH-antigen binding was measured via *k* experimental replicas and *k’* synthesis replicas. These *k x k’* measurements are aggregated in the following way to obtain one normalized log KD measurement:

1. We standardize all experimental replicates so that each experimental replicate *i* has, over all VHHs, mean 0 and standard deviation 1, for each antigen target. We ignore non-binding (Infinity) measurements to compute the mean and standard deviation.
2. We next apply robust normalization so that the parent (VHH-72) binding to each target has median 0 and IQR 1, by subtracting the median parent binding from all VHH binding measurements, then dividing them by the parent binding’s IQR. This operation is done for each *experimental* replica: for each of the *k* experimental replicas, we compute the parent binding’s median and IQR over all *k’* synthesis replicas. Note that this step will maintain non-binding (Infinity) measurements.
3. Finally, we compute the median binding of a candidate VHH to its target (over all synthesis and experimental replicas of the VHH) as the normalized log KD.

We compute *p*-values for round 3 sequences in the following way:

1. We standardize all experimental replicates as in step 1 above.
2. We use a Mann-Whitney *U*-test comparing all bindings of a candidate VHH’s synthesis and experimental replicas to the synthesis and experimental replicas of the parent sequence for sequences with 18 measurements in the last round (we did not compute *p*-values for the earlier, “exploration” rounds).
3. We apply the Bonferroni correction (*α*=0.05) for multiple hypothesis testing.

### Design of the 2nd and 3rd library

For the 2nd and 3rd library we expand the search space to include variants with mutations across all three CDRs by switching to 300nt oligos, and slightly truncating the design region to cover the region from the start of CDR1 to the end of CDR3. This excluded the two FW1 positions that were included in round 1, reducing the design region to the 34 CDR positions plus the 12 FWR positions from round 1 that fall in FWR2 and FWR3. In addition we disallowed mutations to cysteine, and allowed mutations to histidine.

For the 2nd and 3rd library, we used supervised model-based optimization (MBO), which involves three steps:

1. Train a supervised model *f* to predict the binding affinity *f*(*x_i_*) of n different CoV targets j given a nanobody sequence *x_i_*
2. Define an acquisition function *a*(*x_i_*) based on *f*(*x_i_*) and generate a large batch of new candidate sequences by optimizing the acquisition function.
3. Select (a diverse set of) sequences from the generated candidate sequences. Each of the three steps will be described in more detail below

#### Model training

For model training, we subtracted normalized log KD values *y_ij_*between VHH *i* and target *j* from the weakest binding measurement to target *j*. The resulting affinity *y’_ij_ = max_i_ y_ij_ - y_ij_* is such that higher values correspond to stronger binding. We encoded each sequence *x_i_* by concatenating its one-hot representation with ten AAIndex features per amino acid obtained through principal component analysis^36^. To leverage information about non-binding events, we trained classification models *f_c_* to distinguish between non-binding (*y’_ij_* = -∞) and binding (*y’_ij_*> -∞) measurements; these classification models return scalars in [0, 1] representing the probability of binding to each target. We also trained regression models *f_r_* by minimizing the mean-squared error, where we masked out non-binding events represented by an infinitely low affinity (*y’_ij_* = -∞). With the rationale to ignore predictions of the regressor whenever the classifier predicts a sequence as non-binding, we then combined both types of models by multiplying the predicted binding affinities of the regressor by the predicted binding probability of the classifier into a final binding score *f_r_*(*x_i_*) *x f_c_*(*x_i_*).

We used binding data for round 1 sequences against 8 targets to train models with which to design round 2 sequences. These 8 targets include the SARS-CoV1_RBD and SARS-CoV2_RBD wildtype, and all CoV2 mutants that were assayed in round 1 except V483A and R408I, for which binding did not correlate strongly with the other CoV2 targets (Extended Data Fig. 1):

- SARS-CoV1_RBD
- SARS-CoV2_RBD
- SARS-CoV2_RBD_G502D
- SARS-CoV2_RBD_N439K
- SARS-CoV2_RBD_N501D
- SARS-CoV2_RBD_N501F
- SARS-CoV2_RBD_S477N
- SARS-CoV2_RBD_V367F

To train the models used to design the third library, we used experimental measurements collected in both round 1 and round 2 against a total of 12 protein targets. These include all 8 targets listed above, together with additional data collected in round 2 against SARS-CoV1_RBD and SARS-CoV2_RBD and the following 4 targets that were introduced in round 2:

- SARS-CoV2_RBD_N501Y
- SARS-CoV2_RBD_N501Y+K417N+E484K
- SARS-CoV2_RBD_N501Y+K417T+E484K
- SARS-CoV2_RBD_R408I

See Extended Data Table 1 for the list of all CoV variants that were assayed in all three designed libraries, and Extended Data Table 2 for a summary of the VHH sequence variants tested in each library.

We used an ensemble light gradient boosted (LGB) model to design the sequence library that was experimentally tested in round 2, and both an ensemble LGB and ensemble CNN model to design the sequence library that was experimentally tested in round 3. While we trained multiple ensemble LGB models for different output targets j separately, we trained one multi-task CNN model to predict binding for all targets jointly. We chose model hyper-parameters by random five-fold cross-validation, and retrained models with the selected hyper-parameters using the full dataset. These models were then optimized to design sequences for experimental validation as outlined below.

#### Model optimization

After model training, we optimized the model in-silico over multiple generations to generate candidate sequences with high predicted affinities. As an optimization objective, we used the weighted average a(x_i_) = w_j_ y’_ij_ over predictions y’_ij_ of targets j with different target weights w_j_. To primarily optimize binding to CoV-2 while maintaining binding to CoV-1, we assigned a weight of 2/3 to the sum of all CoV-2 targets, i.e., we weighted each of the N CoV-2 targets by w_j_ = 2/3 * 1/N, and weighted the CoV-1 targets by 1/3.

We constrained the number of mutations *d*(*x_i_*) of generated sequences x_i_ relative to VHH-72 by decreasing the objective function *a*(*x_i_*) linearly by max(0, *d*(*x_i_*) - □) * ɣ for different distance thresholds □ and a decay factor ɣ = 0.01, and discarded sequences with more than 15 mutations. In this way, we constrained the optimization within a trust-region around the parent sequence, excluding model predictions for sequences that we felt might be too distant from the training data.

We seeded the optimization with the best 100 sequences discovered so far, and optimized *a*(*x_i_*) using using population-based blackbox optimization (P3BO), which involves an ensemble of discrete blackbox optimization approaches^24^, including greedy local search (SMW), regularized evolution (Evolution)^37^, and design by adaptive sampling^38^. P3BO iteratively optimizes the objective function by sampling candidate sequences from the ensemble of optimization algorithms, where the contribution of each algorithm is weighted by its performance at discovering sequences with a high objective value in the past. This approach enables the discovery of diverse and high-scoring sequences more rapidly than sampling sequences from a single optimization algorithm when working with *in silico* oracles^24^. We performed 100 optimization steps (generations) with 100 sequences per batch. To increase the diversity of designed sequences, we repeated the optimization five times with different random seeds. We further used different different distance thresholds □ (□=5 for library 2, □=6, 7, 8 for library 3) to explore the sequence space at different distances relative to the parent sequence. These choices reflect our prior experience that the region around the parent sequence where the model can be trusted expands as more training data is available^24^. Lastly, we optimized both the LGB and CNN models that were used to design the 3rd library separately.

#### Selection of the 2nd library

To select round 2 sequences, we selected the best (i.e. highest *a*(*x_i_*)) 6800 sequences that only contain mutations in CDR1 or CDR2, the best 6800 sequences with that only contain mutations in CDR3, and the best 14000 sequences with mutations in all three CDR region. Additionally, we selected the best 700 sequences that only contain mutations at positions near the binding domain (IMGT number 58, 66, 112B, 112A, 113, 114). We discarded sequence duplicates.

#### Selection of the 3rd library

To select round 3 sequences, we first pre-selected the best sequences with a reward above the 75th percentile for each model after deduplicating sequences. To increase the diversity of the selected sequences, we then clustered the pre-selected sequences of both the CNN and LGB model separately into eight clusters (16=2×8 clusters in total) using the K-means algorithm and Euclidean distance between AAIndex-featurized sequences as distance metric. Finally, we selected the best 4000 sequences of the CNN and LGB model equally from each of the eight clusters per model, resulting in 8000 total sequences.

### Recombination baseline for validating ML-guided sequence design

For the first library, we regard the set of all permitted single mutant sequences as a baseline library, since this type of deep mutational scanning approach is often used in protein engineering and antibody discovery. For round 2, we built a number of baselines, described in detail below. To identify the best sequences from round 1, we considered all sequences as candidates whether designed using the round 1 baseline strategy (single mutants of VHH-72) or the round 1 ML-guided strategy (VAE). As noted above, due to synthesis constraints all sequence variants tested in the first round contained mutations in either CDRs 1+2 or CDR 3. For the second and third round, the synthesis constraints were changed so that the entire region from the start of CDR1 to the end of CDR3 was available for modification.

Singles of CDR12 mutants: We identified the best 100 sequences from round 1 with mutations in CDRs 1+2. For each of these 100 founder sequences, we then drew 12 single mutants at random from the best 200 single mutant variants measured in round 1, and added each single mutant to a separate copy of the founder sequence. In the event of a collision (where a best single mutant is already contained in a founder sequence) we added an additional single mutant to yield a total of 2400 new sequence designs.

Singles of CDR3 mutants: We identified the best 100 sequences from round 1 with mutations in CDR 3. For each of these 100 founder sequences, we then drew 12 single mutants at random from the best 200 single mutant variants measured in round 1, and added each single mutant to a separate copy of the founder sequence. In the event of a collision (where a best single mutant is already contained in a founder sequence) we added an additional single mutant to yield a total of 2,400 new sequence designs.

Recombinations: We used a simple additive model with the measurements from round 1 to identify the 300 best combinations of (i) a sequence variant with mutation(s) in CDRs 1+2 and (ii) a sequence variant with mutation(s) in CDR 3.

Singles of recombinations: Using the 300 recombinants described above, we treated each as a founder sequence and drew 8 single mutants at random from the best 200 single mutant variants measured in round 1, adding each single mutant to a separate copy of the founder sequence (again dropping down the list of singles in case of any collisions) to yield an additional 2,400 new sequence designs.

Shuffled: We randomly shuffled the amino acids within each CDR to provide an additional very simple baseline.

For round 3, we reprised both of the baseline strategies used in round 2. We first selected a set of best starting sequences from all of the round 1 and the round 2 baseline sets, by (i) assigning non-binders the weakest binding measurement from each round, (ii) averaging values over replicas and then over CoV2 targets, (iii) removing sequences with mutations outside the set that could be reached by synthesis for round 3, (iv) removing all sequences with values significantly worse than parent. These steps result in a set of 16 best singles from round 1, and 148 best sequences drawn from round 1 (9 VAE-designed sequences) and the round 2 baseline (139 sequences) as starting points.

We then constructed two sets of baseline sequences. The first added each best single to each of the best 148 sequences and provided a score for each combination by assuming a simple additive model. From this large set, we chose the 250 sequences with the highest predicted scores, and in addition 250 other sequences sampled from the remaining set at random proportional to their predicted scores. For the second set, we combined the mutations from each of the best 148 sequences with each other, again assuming a simple additive model. From this set we chose the 500 with highest predicted score and in addition another 500 sequences sampled from the remaining set at random proportional to their predicted scores.

### High throughput experimental (Alpha Seq) assay

#### Coronavirus strains

The following Coronavirus strains were used in this study: SARS-CoV-2, isolate Wuhan-Hu-1, Genbank MN908947; SARS-CoV-1 Urbani, Genbank AY278741; Middle East respiratory syndrome-related coronavirus, NCBI Reference Sequence YP_009047204.1; BM48-31, NCBI Reference Sequence YP_003858584.1; ZXC21, GenBank AVP78042.1; RaTG13, GenBank QHR63300.2; Rf1 GenBank ABD75323.1; HKU3-1, UniProtKB/Swiss-Prot Q3LZX1.1; LYRa11, GenBank AHX37558.1; ZC45, GenBank AVP78031.1; WIV1, GenBank AGZ48828.1; and Rp3, UniProtKB/Swiss-Prot Q3I5J5.1.

#### Yeast media

Yeast peptone dextrose (YPD), yeast peptone galactose (YPG), and synthetic drop out (SDO) media supplemented with 80 mg/L adenine were made according to standard protocols. Suppliers used for our yeast media are as follows: Bacto Yeast Extract (Life Technologies), Bacto Tryptone (Fisher BioReagents), Dextrose (Fisher Chemical), Galactose (Millipore Sigma), Adenine (ACROS Organics), Yeast Nitrogen Base w/o Amino Acids (Thermo Scientific), SC-His-Leu-Lys-Trp-Ura Powder (Sunrise Science Products), L-Histidine (Fisher BioReagents), L-Tryptophan (Fisher BioReagents), Uracil (ACROS Organics), and Bacto Agar (Fisher BioReagents).

#### Isogenic yeast plasmid transformation

AlphaSeq compatible plasmids encoding yeast surface display cassettes were constructed by Twist Bioscience and resuspended at 100ng/µL. 100ng of plasmid was digested with PmeI enzyme for 1hr at 37°C to linearize, leaving chromosomal homology for integration into the ARS314 locus at both the 5’ and 3’ ends as described in (*Younger PNAS*). Yeast transformations were performed with Frozen-EZ Yeast Transformation Kit II (Zymo Research) according to manufacturer’s instructions. Yeast were plated on SDO-Trp plates and grown at 30°C for 2-3 days. Successful transformants were struck out onto yeast peptone adenine dextrose (YPAD) plates and grown overnight at 30°C.

#### Protein expression validation – Flow cytometry

Yeast was inoculated in YPAD and grown overnight at 30°C. Yeast were labeled with FITC-anti-C-myc antibody (Immunology Consultants Laboratory, Inc.) in PBS (Gibco) + 0.2% BSA (Thermo) for 30 minutes at RT. Yeast were pelleted and resuspended in PBS + 0.2% BSA and read on a LSRII cytometer.

#### Target library DNA library construction

AlphaSeq compatible fragments were synthesized by Twist Bioscience and were resuspended at 1 ng/µL in molecular grade water and pooled together. Fragment libraries were PCR amplified using KAPA DNA polymerase (Roche). A second DNA fragment with a randomized DNA barcode was PCR amplified. Fragments were run on a 0.8% agarose gel and extracted using Monarch Gel Purification kit (NEB).

#### VHH library DNA construction

200bp or 300bp Oligo Pools were synthesized by Twist Bioscience and resuspended at 20 ng/µL in molecular grade water. Oligo fragments were PCR amplified using KAPA DNA polymerase (Roche). qPCR was terminated before saturation to minimize PCR bias, generally between 8-15 cycles. Upstream and downstream fragments which contained homology to the Oligo fragments were PCR amplified using KAPA DNA polymerase (Roche). A 3 piece Gibson assembly reaction using Gibson Assembly Master Mix (NEB) was performed following the manufacturer’s instructions. The Gibson assembly reaction was cleaned up using KAPA beads (Roche) and the entire reaction was used as a template to PCR amplify a fully assembled AlphaSeq compatible fragment using KAPA DNA polymerase (Roche). qPCR was terminated before saturation to minimize PCR bias, generally between 12-15 cycles. A second DNA fragment with a randomized DNA barcode was PCR amplified. Fragments were run on a 0.8% agarose gel and extracted using Monarch Gel Purification kit (NEB).

#### Yeast library transformation

MATa or MATalpha AlphaSeq yeast were grown for 6 hours at 30°C in YPAG media to induce SceI expression, as described in (*Younger PNAS*). All spin steps were performed at 3000 RPM for 5 minutes. Yeast were spun down and washed once in 50 mL 1M Sorbitol (Teknova) + 1mM CaCl_2_ solution. Washed yeast were resuspended in a solution of 0.1M LiOAc/1mM DTT and incubated shaking at 30°C for 30 minutes. After 30 minutes, yeast was spun down and washed once in 50 mL 1M Sorbitol + 1mM CaCl_2_ solution. Yeast was resuspended to a final volume of 400 µL in 1M Sorbitol + 1mM CaCl_2_ solution and incubated with DNA for at least 5 minutes on ice. Yeast were electroporated at 2.5kV and 25 uF (BioRad). Immediately following electroporation, yeast were resuspended in 5 mL of 1:1 solution of 1M Sorbitol:YPAD and incubated shaking at 30°C for 30 minutes. Recovered yeast cells were spun down and re-suspend in 50 mL of SDO-Trp media and transferred to a 250mL baffled flask. 20 µL of resuspended cells were plated on SDO-Trp to determine transformation efficiency. Both the flask and plate were incubated at 30°C for 2-3 days. After 2-3 days, transformation efficiency was determined by counting colonies on the SDO-Trp plate.

#### Nanopore barcode mapping

Genomic DNA from yeast libraries was extracted using Yeast DNA Extraction Kit (Thermo Scientific) following the manufacturer’s instructions. A single round of qPCR was performed to amplify a fragment pool from the genomic DNA containing the gene through the associated DNA barcode. qPCR was terminated before saturation to minimize PCR bias, generally between 15-20 cycles. The final amplified fragment was concentrated with KAPA beads, quantified with a Quantus (Promega), prepped with either a SQK-LSK-109 or SQK-LSK-110 ligation kit (Oxford Nanopore) and sequenced with either a Minion R9 or R10 flow cell (Oxford Nanopore) following the manufacturer’s instructions.

#### Library-on-Library AlphaSeq Assays

2 mL of saturated MATa and MATalpha library were combined in 800 mL of YPAD media and incubated at 30°C in a shaking incubator. 3 technical replicates were performed for each assay. After 16hr, 100 mL of yeast culture was washed once in 50 mL of sterile water and transferred to 600 mL of SDO-lys-leu with 100 nM ß-estradiol (Sigma) for 24hr. After 24hr, 100 mL of yeast was transferred to fresh SDO-lys-leu with 100 nM ß-estradiol for an additional 24hr.

#### Library preparation for Next-Generation Sequencing

Genomic DNA was extracted using Yeast DNA Extraction Kit (Thermo Scientific) following manufacturer’s instructions. qPCR was performed to amplify a fragment pool from the genomic DNA and to add standard Illumina sequencing adaptors and assay specific index barcodes. qPCR was terminated before saturation to minimize PCR bias, generally between 23-27 cycles. The final amplified fragment was concentrated with KAPA beads, quantified with a Quantus (Promega), and sequenced with an NextSeq 500 sequencer (Illumina).

#### Dissociation constant estimation

Both diploid barcode pairs counts, and haploid barcode counts were generated using Illumina sequencing as described above. Counts of diploid barcode pairs were normalized by dividing them by the product of each corresponding haploid barcode counts to account for differences in library representation. A standard curve of proteins with known interacting partners and dissociation constant values was included in the assay and used to extrapolate dissociation constant values for the experimental protein interactions.

### Validation experiments

#### Sequence selection for BLI and Pseudovirus neutralization assays

To confirm the high throughput results measured by the AlphaSeq assay we also measured binding of VHH-72, the best single-mutant sequence of the 1st library (Seq12), and 19 designed sequences using Biolayer Interferometry (BLI). We selected 19 designed sequences by first excluding sequences that bound to negative control targets, an indication of unspecific binding, and sequences with mutations to cysteine (allowed in round 1 only). From the set of both baseline and ML-designed sequences, we then pre-selected the 200 sequences with 1) the smallest Benjamini-Yakutelli adjusted p-value, 2) the smallest median-aggregated affinity, and 3) the smallest mean-aggregate affinity, and took the intersection of these sets. To increase diversity, we then k-means clustered the resulting 61 sequences into 10 clusters, and selected from each cluster the best one or two sequences with the smallest p-value to achieve a total of 19 sequences. These 19 sequences were tested for protein expression and using BLI for binding to the SARS-CoV-1 and SARS-CoV-2 RBDs (Extended Data Fig. 5).

Of these 19 sequences, we found that 13 sequences (MLp-designed Seqs 1-11, VHH-72 and Seq 12) showed high expression (>10 mg/L) in *E. coli.* These 13 high expression sequences were subsequently tested in pseudovirus neutralization assays against SARS-CoV2-WT (Extended Data Fig. 5), with the ML-designed sequences achieving up to 200-fold improved neutralization over VHH-72. Furthermore these 13 high expression sequences were also tested in additional BLI and pseudovirus neutralization assays against the SARS-CoV-1 WT, the SARS-CoV-2 WT, and the Delta, and Omicron variants; results are reported in Figure 3 and the main text.

#### Protein expression and purification

Protein expression constructs were cloned into a modified pET28 b(+) vector containing a His-tag and an ampicillin resistance marker. Sequence verified plasmids were transformed into BL21(DE3) *E. coli* cells (New England Biolabs) and plated on LB medium supplemented with 50 μg mL^−1^ ampicillin. Protein constructs were expressed using autoinduction protocol described previously [doi: 10.1016/j.pep.2005.01.016]. Briefly, 100 ml of auto-induction media containing ampicillin was inoculated with a single transformed colony, shaken at 37°C for 8 h, followed by 19°C for 36 hours. Cultures were pelleted and stored at −20°C until purification. Pellets were thawed on ice and resuspended in 30 ml lysis buffer (50 mM Tris pH 8.0, 500 mM NaCl, supplemented with 1x pierce protease inhibitor tablets and 1 mM PMSF). Cells were lysed by high pressure homogenization at 12,000 psi. The lysates were clarified by centrifugation at 18,500*g* at 4°C for 30 minutes. To purify recombinant protein, clarified lysates were passed through 5 ml HisTrap columns (Cytiva) equilibrated with 50mM Tris pH 8.0, 500 mM NaCl and 20 mM imidazole on Akta Pure FPLC system and UNICORN software (Cytiva). HisTrap columns were washed with 10 CVs of wash buffer. Bound protein was eluted using gradient flow of 5 CVs elution buffer at 10% - 100% gradient slope. Fractions containing the eluted protein were assessed by SDS-PAGE gel. Samples were separated on Bis-Tris NuPAGE 4 to 12% gels (Thermo Scientific), followed by Coomassie Blue gel staining. Fractions were pooled, concentrated, and further purified using a HiLoad 16/600 Superdex 200 pg column (Cytiva) on AKTA Pure FPLC equilibrated with 1X PBS. Purity and final analysis were performed using SDS-PAGE gel under non-reducing and reducing conditions.

#### Kinetic binding measurements

Biolayer interferometry (BLI) was used to measure dissociation constants for each VHH-target pair. RBD variants were purchased commercially (Sino Biological) and biotinylated with an EZ-Link NHS-PEG4 biotinylation kit (Thermo Scientific) according to the manufacturer’s instructions. Biotinylated target proteins were loaded onto SA or SAX biosensors (Sartorius) of an Octet Red96e (Sartorius) until a 1 nm shift was reached. Following a 120 s baseline step, binding of each VHH was measured using a 60 s association phase and a 180 s dissociation phase. A sample with no VHH was used for background subtraction and normalization. Three concentrations of each VHH was assayed, and a global fit was applied using the Octet Analysis Studio (v12.2.2.26, Sartorius) to calculate association and dissociation rates.

#### Pseudovirus neutralization assays

Lentiviral particles pseudotyped with SARS-CoV-2 spike proteins were produced and titered as described previously^39^ with minor modifications. 293T cells were cultured in Dulbecco’s Modified Eagle Medium (DMEM) supplemented with 10% heat-inactivated fetal bovine serum and 100 U/mL penicillin-streptomycin (Gibco). The cells were grown in a high-humidity incubator at 37°C and 5% CO_2_. Approximately 1 x 10^7 293T cells were seeded in 10 ml of DMEM per T75 flask and incubated for 16-24 hours until cells were 50-70% confluent. The cells were transfected using Lipofectamine 2000 transfection reagent (Thermo Scientific) according to the manufacturer’s protocol with the following plasmid mix per T75 flask: 7.8 mg of lentiviral backbone: Luciferase-IRES-ZsGreen (BEI NR-52516); 1.716 mg each of helper plasmids: HDM-Hgpm2 (BEI NR-52517), pRC-CMV-Rev1b (BEI NR-52519), and HDM-tat1b (BEI NR-52518); and 2.65 mg viral entry protein: SARS-CoV-2 Spike, SARS-CoV-2 Delta Spike variant, SARS-CoV-2 Omicron Spike variant, or SARS-CoV-1 Spike with 21 amino acids deleted from the C-terminus. At 4 hours post-transfection, the media was removed and replaced with 15 ml fresh, prewarmed DMEM. At 48-60 hours post-transfection, the pseudovirus was harvested by collecting the supernatant and filtering through a 0.22 mm filter. Pseudovirus aliquots were stored at -80°C. The pseudovirus was later titered in order to determine the amount used in the neutralization assay. 293T-ACE2 cells (gift of the Bloom lab) were cultured similarly to 293T cells. 96-well flat-bottom polystyrene plates with opaque walls were coated with Poly-L-Lysine (Millipore Sigma) according to the manufacturer’s protocol to improve cell adherence. Coated 96-well plates were seeded with 1.25 x 10^4 293T-ACE2 cells per well in 100 µl of media and incubated for 16-24 hours. Pseudovirus aliquots were thawed on ice.. The VHH samples were serial diluted in a separate 96-well plate, mixed with pseudovirus, and incubated at 37°C for one hour. Culture media was carefully removed from the plate of 293T-ACE2 cells and 100 µl of VHH/pseudovirus mixture was added on top of the cells. Polybrene (Sigma-Aldrich) was added to a final concentration of 5 mg/ml. At 48-60 hours post-infection, luminescence was measured using the Bright-Glo Luciferase Assay System (Promega). Data was analyzed and plotted with Prism. Virus infectivity was calculated by normalizing the cells-only condition to 0% and the virus+cells-only condition to 100%.

### Structural analysis

We built the structural model shown in Figure 4 by aligning the RBD domain of CoV-2 (PDB 6m17), with the RBD domain of CoV-1 combination with VHH72 (PDB 6waq). We further aligned the RBD domain of CoV-2 Delta (PDB 7w9c), and the RBD domain of Omicron (PDB 7wpb), with the RBD domain of CoV-2 (PDB 6m17), to the highlight the mutations of Delta and Omicron, respectively. We modeled the mutations of VHH1 shown in yellow sticks by using the PyMOL mutagenesis tools and selecting rotamers with the lowest energy conformation. All analyses were performed with PyMOL 2.5.4. We analyzed polar contacts such as those shown in Figure 5C between N370 and G55 using the PyMol command *distance* with cutoff=3.2 and mode=2 (polar contacts only).

## Data Availability Statement

The experimental data of all three design libraries, model checkpoints, and evaluation data to reproduce results will be made publicly available upon submission of the manuscript. This dataset also includes literature sequences and DBR sequences that are not described in this manuscript. Furthermore, the code required to synthesize, process and analyze the experimental data will be provided for download, together with the iPython notebooks that reproduce the analysis figures from the main text. Further details available from the authors upon request.ee

## Code Availability Statement

Scikit-learn, Tensorflow, and LightGBM where to implement and train all models using the architectures described in Methods. The training and validation datasets used to build each model are available as part of the experimental dataset released as described in Data Availability. We provide a checkpoint of the trained model used to design VHH sequences, together with code that loads the model and enables the user to run inference for candidate VHH sequences of their choice.

## Author contributions

J.R, M.F., L.J.C., R.L., and D.Y. conceived the study. C.A., Z.M., Z.R.M., B.A., and L.J.C. analyzed data and designed, implemented and used machine learning models to design sequences. B.J., E.E., R.E., C.L., C.S., D.G., J.N., M.K., M.M., P.S., C.C., S.H., S.S., K.A., S.E., A.M., S.O., and K.H. performed wet-lab experiments and prepared the data. C.A, Z.M., L.J.C., B.J., E.E., R.E., R.L., and D.Y. wrote the manuscript with the input of all authors.

## Acknowledgements

We thank David Belanger and Max Bileschi for countless conversations and guidance throughout this project. We thank the entire A-AlphaBio and Lumen team for their support and guidance throughout the project.

## Extended Data Figures

**Extended Data Fig. 1.**
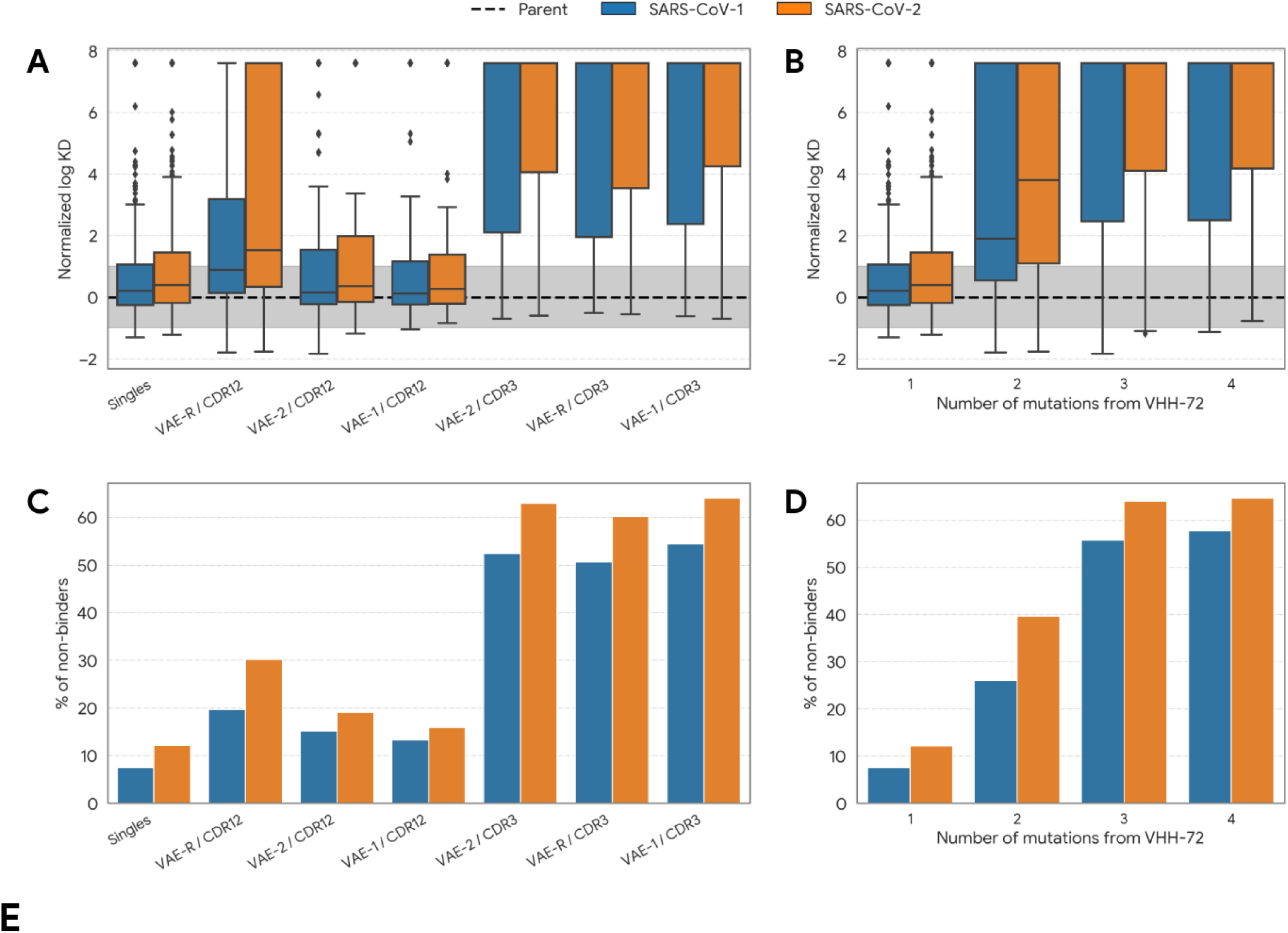
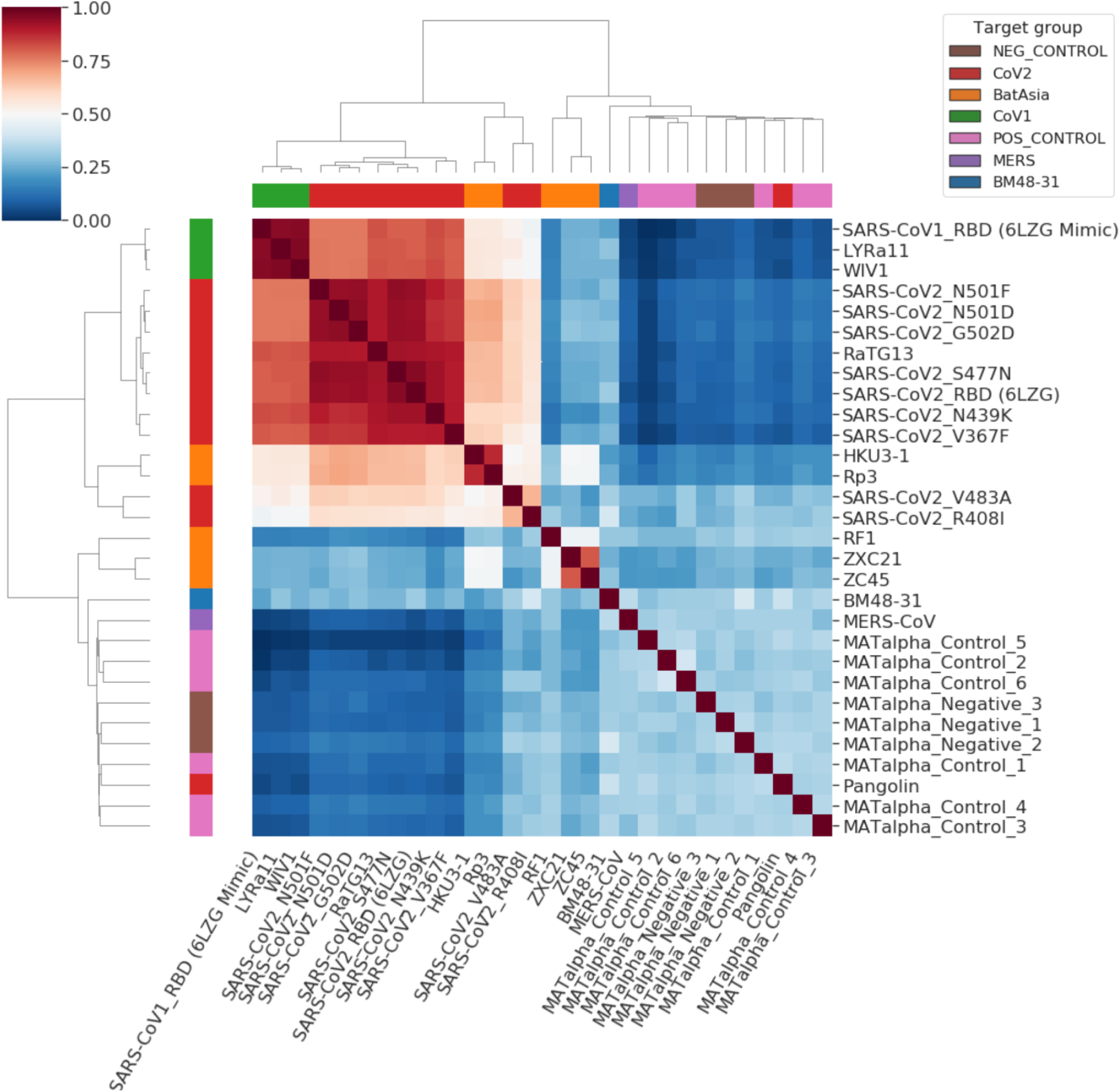
Detailed first library results. (A) Normalized log KD of designed VHHs against SARS-CoV-2 depending on the design strategy. VAE-1 and VAE-2 means that sequences were designed by maximizing the likelihood of the 1st or 2nd VAE model, respectively. VAE-R means that sequences were designed by sampling at random from all sequences that had VAE likelihood above a threshold. Mutating CDR3 has a higher (deleterious) impact on binding, emphasizing the importance of this region in driving binding to SARS-CoV-2. Sequences sampled at random above a likelihood threshold (VAE-R) were less likely to bind tightly than those sampled by maximizing the likelihood of the 1st or 2nd VAE model. (B) Normalized log KD depending on the number of mutations. Since only unsupervised data (OAS sequences) were used to train VAEs, we hypothesize that optimizing these models tends to produce sequences that lose specificity to the SARS-CoV targets at higher number of mutations. (C) Percentage of non-binding sequences by design method. Mutating CDR3 following the VAE scores has the highest likelihood (up to 60%) of designing non-binders, in-line with the conclusions from subfigure A. (D) Fraction of non-binders as a function of the number of mutations away from VHH-72. Because sequences with ≥2 mutants were designed by VAE models that do not consider specificity to either of the SARS-CoV targets, these sequences are more likely to be non-binders. (E) Hierarchical clustering of all CoV and CoV-related targets included in the 1st library based on their AlphaSeq binding affinities. Heatmap represents Spearman’s R between target pairs. Rows and columns are colored by target groups. CoV-2 targets cluster closely, except for SARS-CoV2_V483A and SARS-CoV2-R408I. The latter two outliers were therefore not used for training the ML model to design the 2nd library.

**Extended Data Fig. 2.**
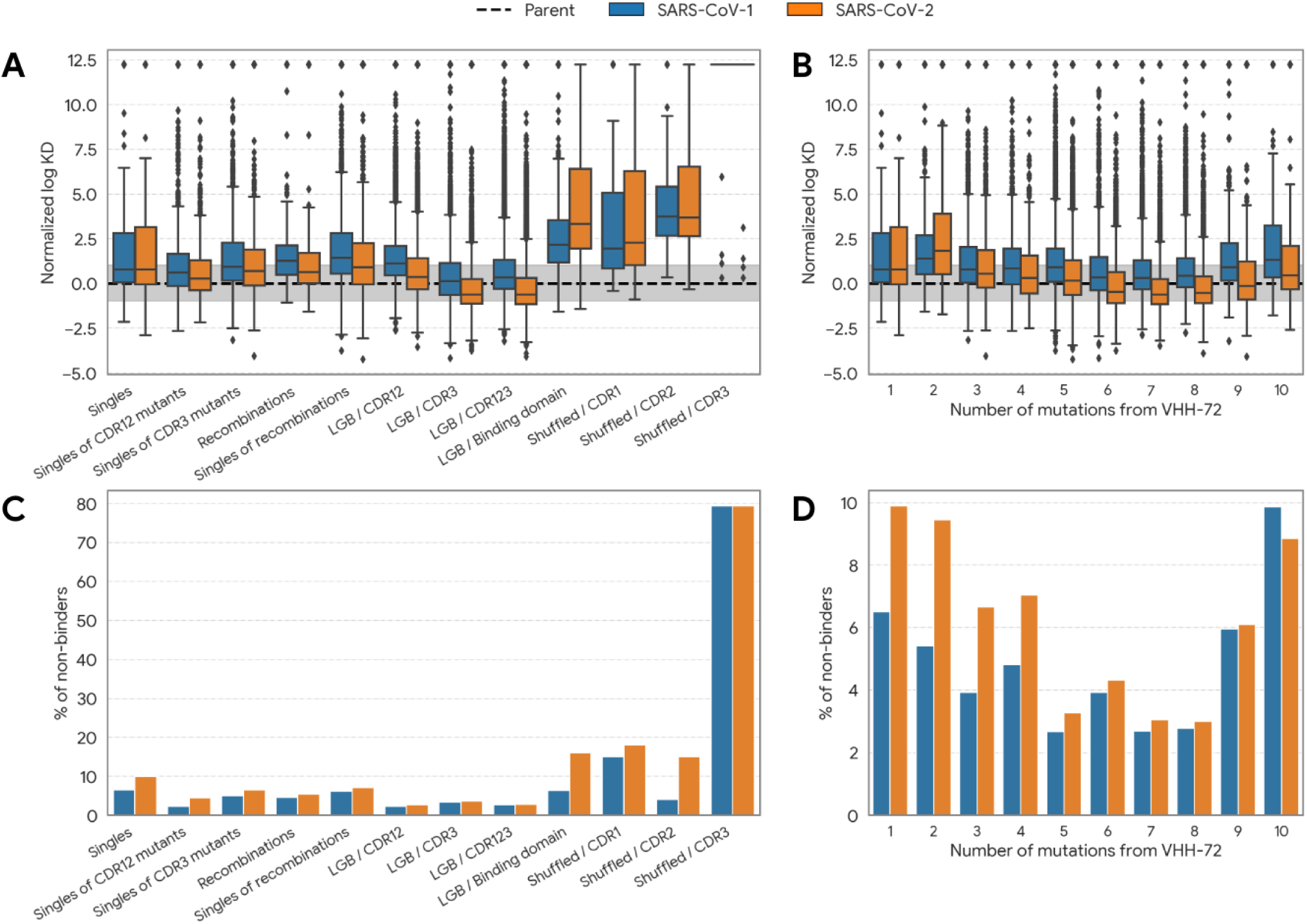
Detailed second library results (sequences duplicated from the first library are not included). (A) Normalized log KD of designed VHHs against SARS-CoV-2 depending on the design strategy. Unlike round 1, model-designed sequences of round 2 improved binding on average. For baseline strategies that modify previous best sequences, recombining sequences tends to perform worse than single mutations. ML-designed sequences use the LGB (Light Gradient Boosted) combined models. Where in the first library, mutating CDR3 was overall deleterious, we now see a marked improvement in sequences that optimize CDR3 over CDRs 1 and 2. Optimizing only those positions thought to make contact with the receptor generally decreased binding by some margin. Finally, shuffled sequences generally reported a large decrease in binding. (B) Normalized log KD as a function of the number of mutations. Models trained using the data from round 1 designed sequences that had improved binding activity over VHH-72 even when several mutations were introduced. Binding strengthens as the number of mutations increases to 6-8 mutations, then decreases as additional mutations are added. We omit shuffled sequences from this plot due to their poor performance (subfigure A). (C) Percentage of non-binders per design strategy. ML-designed sequences that optimize the CDRs design the highest fraction of binding sequences. However, optimizing the binding domain with ML is more likely to ablate binding. Shuffling CDRs is, as expected, deleterious to binding, with CDR3 being particularly sensitive to shuffling. (D) Percentage of non-binders as a function of the number of mutations; as in subfigure B, we removed shuffled sequences from this analysis. Where the previous library saw the number of non-binders increase as a function of the number of mutations, this is no longer the case. Up to 8 mutations away, the number of non-binders to SARS-CoV-2 is lower than 4%. Increasing the number of mutations even further does however increase the fraction of non-binders, as the designed sequences mutate further away from the available training data.

**Extended Data Fig. 3.**
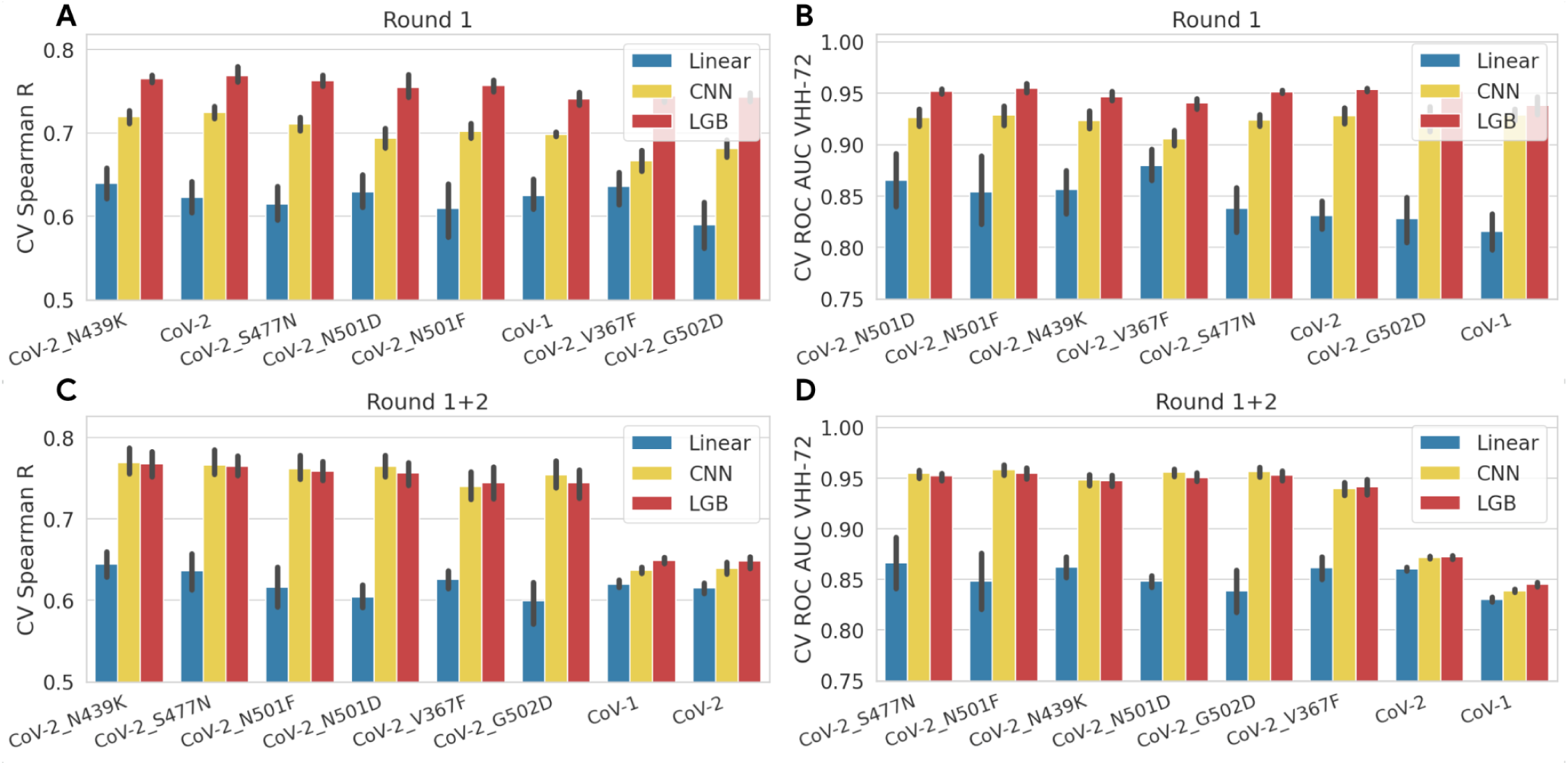
Model cross-validation performance per target. (A) 5-fold cross-validation performance quantified by Spearman’s R of models trained on round 1, for all 8 targets that were assayed in round 1 and round 2. (B) ROC AUC score of model trained on round 1 to correctly classify sequences that bind more tightly than VHH-72. (C-D) Hold-out Spearman’s R and ROC AUC score per target of models trained on both sequences of round 1 and round 2. The plot shows that the LGB and CNN models perform best across all targets. In subfigures A-B the cross-validation set includes data from just the first library, while in C-D, the cross-validation set includes data from both the first and second libraries. For the CoV-1 and CoV-2 WT targets, the model-designed sequences tested in the second library had notably better performance. As a result, their measured affinities were concentrated at affinities similar to, or tighter than that of the VHH-72 parent sequence. This data shift makes the task of ranking held-out sequences or classifying whether they bind more tightly than VHH-72 more difficult, resulting in the performance drop observed in subfigures C-D for the CoV-1 and CoV-2 WT targets.

**Extended Data Fig. 4.**
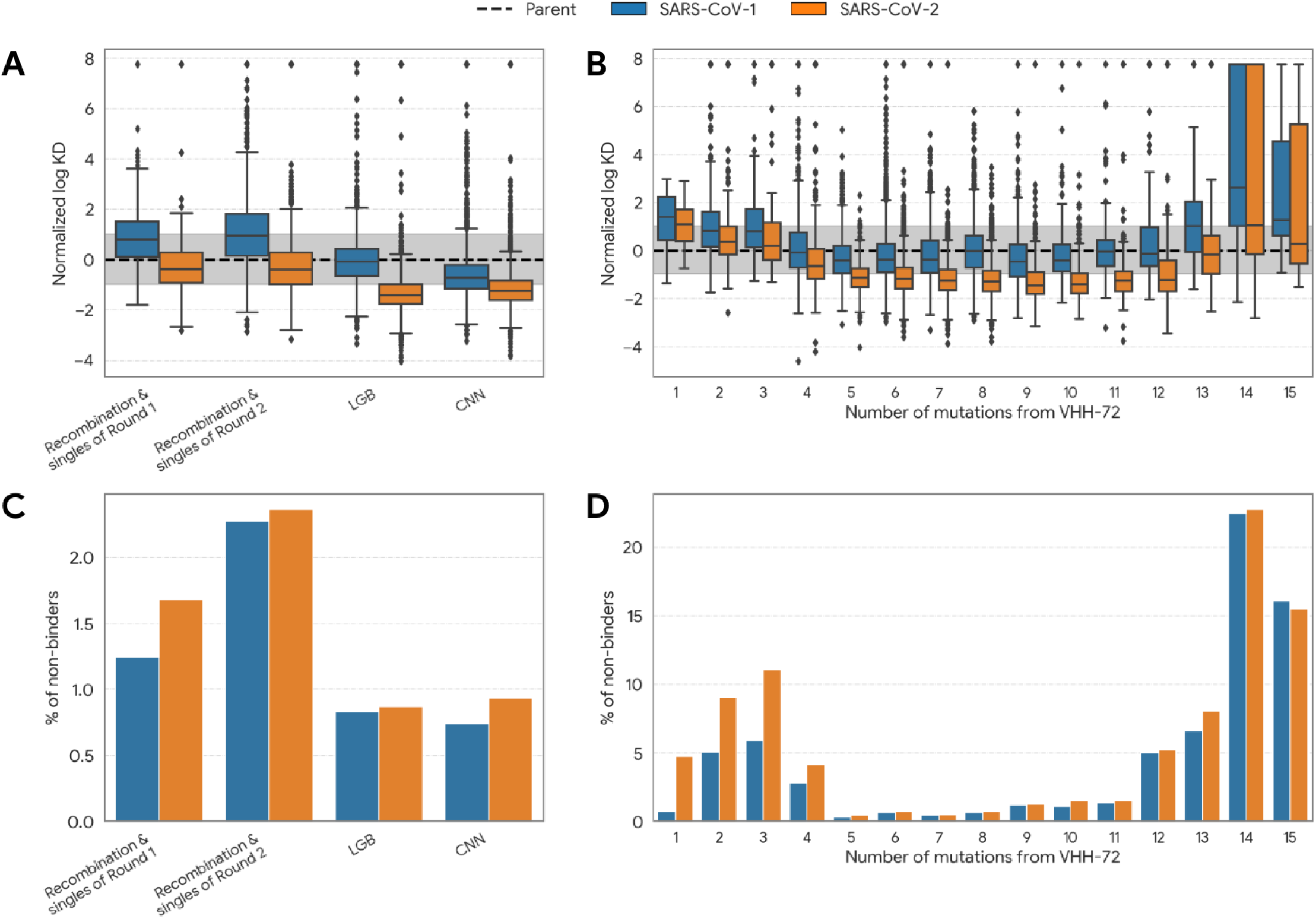
Detailed third library results (sequences duplicated from earlier rounds are not included). (A) Normalized log KD of designed VHHs against SARS-CoV-2 as a function of the detailed model strategy. The average binding to SARS-CoV-2 is stronger than to SARS-CoV-1 across all design strategies. Both types of ML-designed sequences, which use either an LGB or CNN model, achieve stronger bindings than the baseline approaches that recombine and mutate either round 1 or round 2 sequences. (B) Normalized log KD as a function of the number of mutations. Following the trend of the previous round, we are able to move even further away from VHH-72, achieving strong binding to SARS-CoV-2 up to 11 mutations away; however, moving much further dramatically reduces binding. (C) Percentage of non-binders per design strategy. ML-designed sequences are more likely to design binding sequences, but all methods design less than 3% of non-binders. (D) Percentage of non-binders as a function of the number of mutations. In line with the conclusions from subfigure B, designed sequences maintain binding to both SARS-CoV targets up to 11 mutations away; sequences with very few (≤4) or too many (≥12) mutations have a much higher fraction of non-binding sequences.

**Extended Data Fig. 5.**
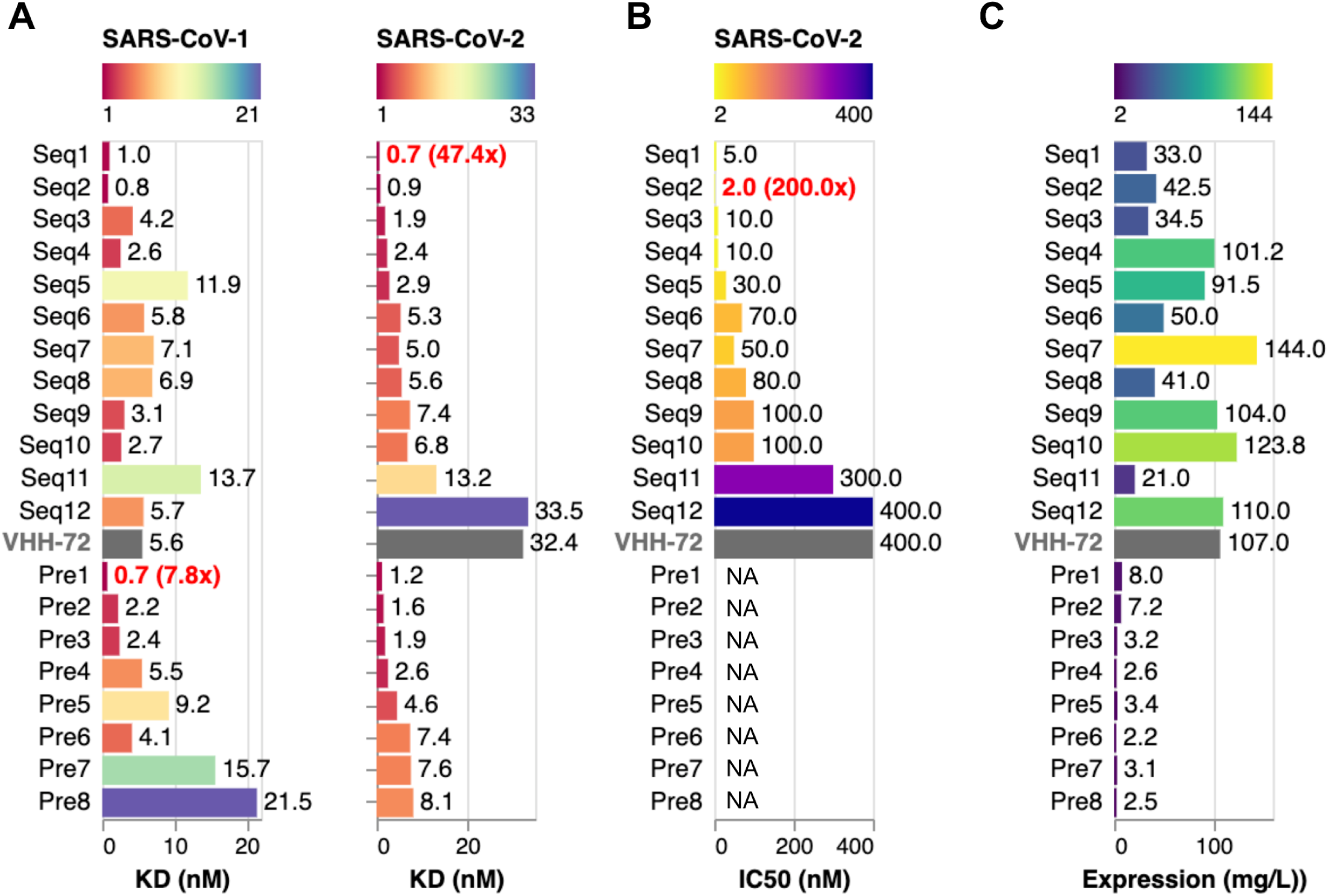
Measurements of the preliminary screening experiment. (A) biolayer interferometry (BLI), (B) pseudovirus neutralization, and (C) protein expression measurements of the 21 sequences (VHH-72, Seq 12 and 19 representative best ML-designed sequences) that were included in the preliminary screening experiment. Pre1-Pre8 denotes sequences with a protein expression level of below 10 mg/L, for which pseudovirus neutralization was not measured, indicated by NA in (B), and which were not included in the follow-up experiment shown in Figure 4.

**Extended Data Fig. 6.**
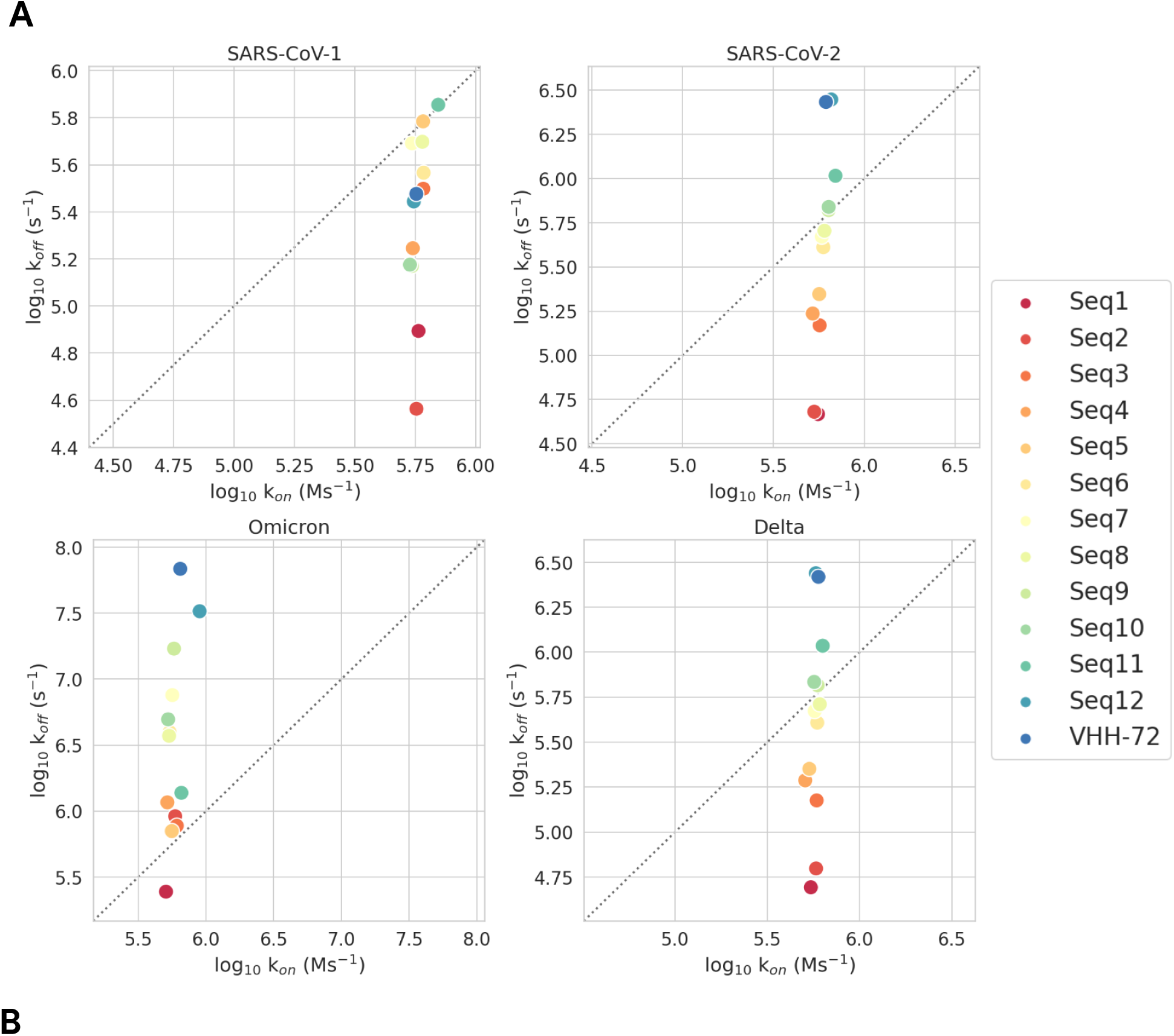
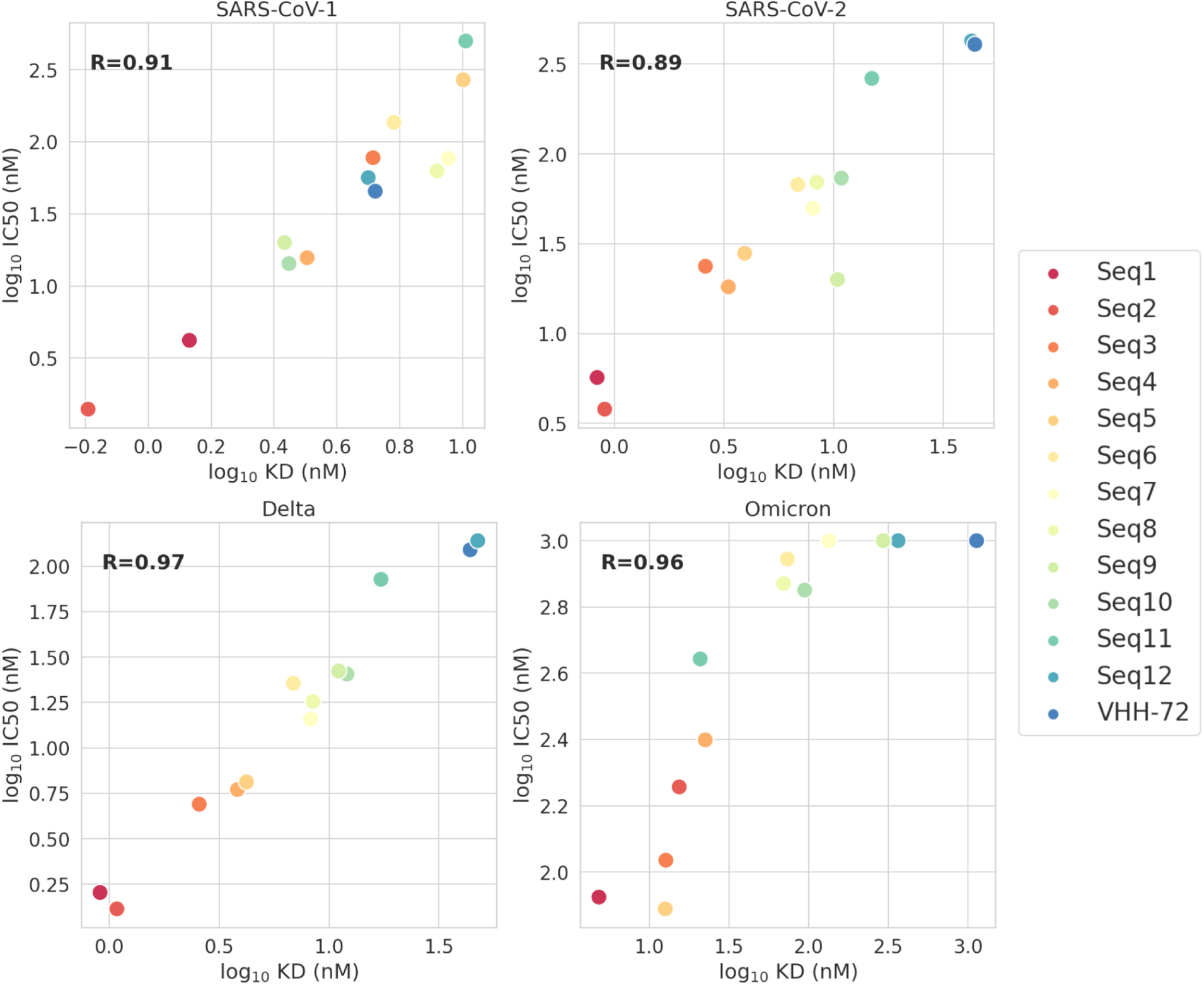
(A) BLI association (k_on_) and dissociation (k_off_) rates. Binding improvements of designed sequences are primarily driven by smaller dissociation rates, while association rates remain similar to VHH-72. (B) Correlation of BLI binding strength and neutralization for SARS-CoV-1, SARS-CoV-2, Delta, and Omicron.

**Extended Data Fig. 7.**
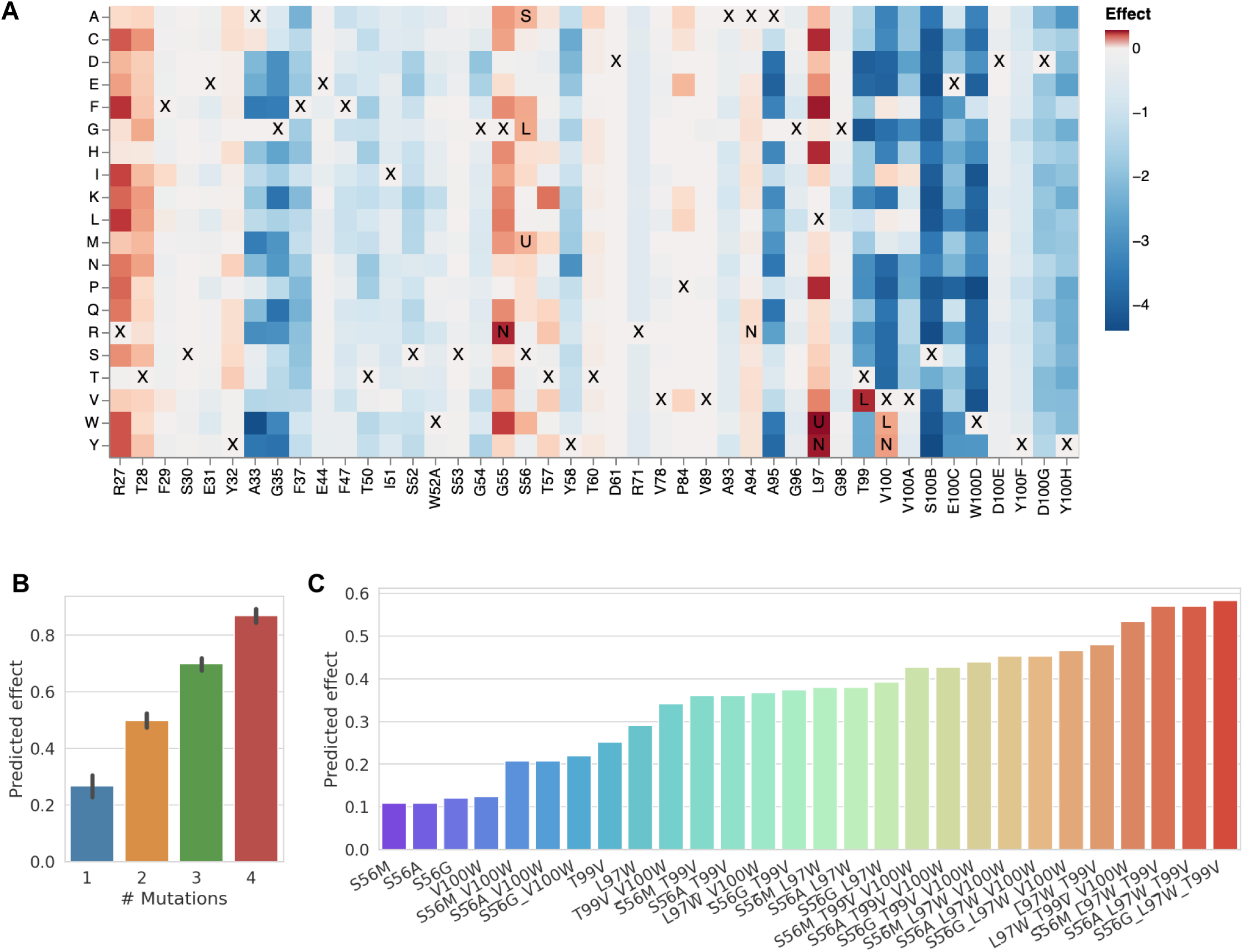
Predicted effect of single-mutants and multi-mutants. (A) Predicted change in binding affinity for all single-mutant variants of VHH-72. Values above and below zero indicate an increase and decrease in binding affinity, respectively. ‘X’ highlights the parent residue of VHH-72. ‘S’, ‘L’, and U’ denotes mutations that were found to improve binding by Schepens et al, Laroche et al, and Sulea et al, respectively. ‘N’ denotes previously undescribed mutations that are most prevalent in our ML designed sequences (Figure 5B). (B) Histogram of the predicted change in binding affinity of all single, double, triple, and quadruple variants of the most prevalent mutations described in the main text (S56G, S56M, S56A, L97W, T99V, V100W). (C) Predicted change in binding of single, double, and triplet variants; overall our model predicts that combining the most frequent single-mutants into higher-order variants will increase binding.

**Extended Data Fig. 8.**
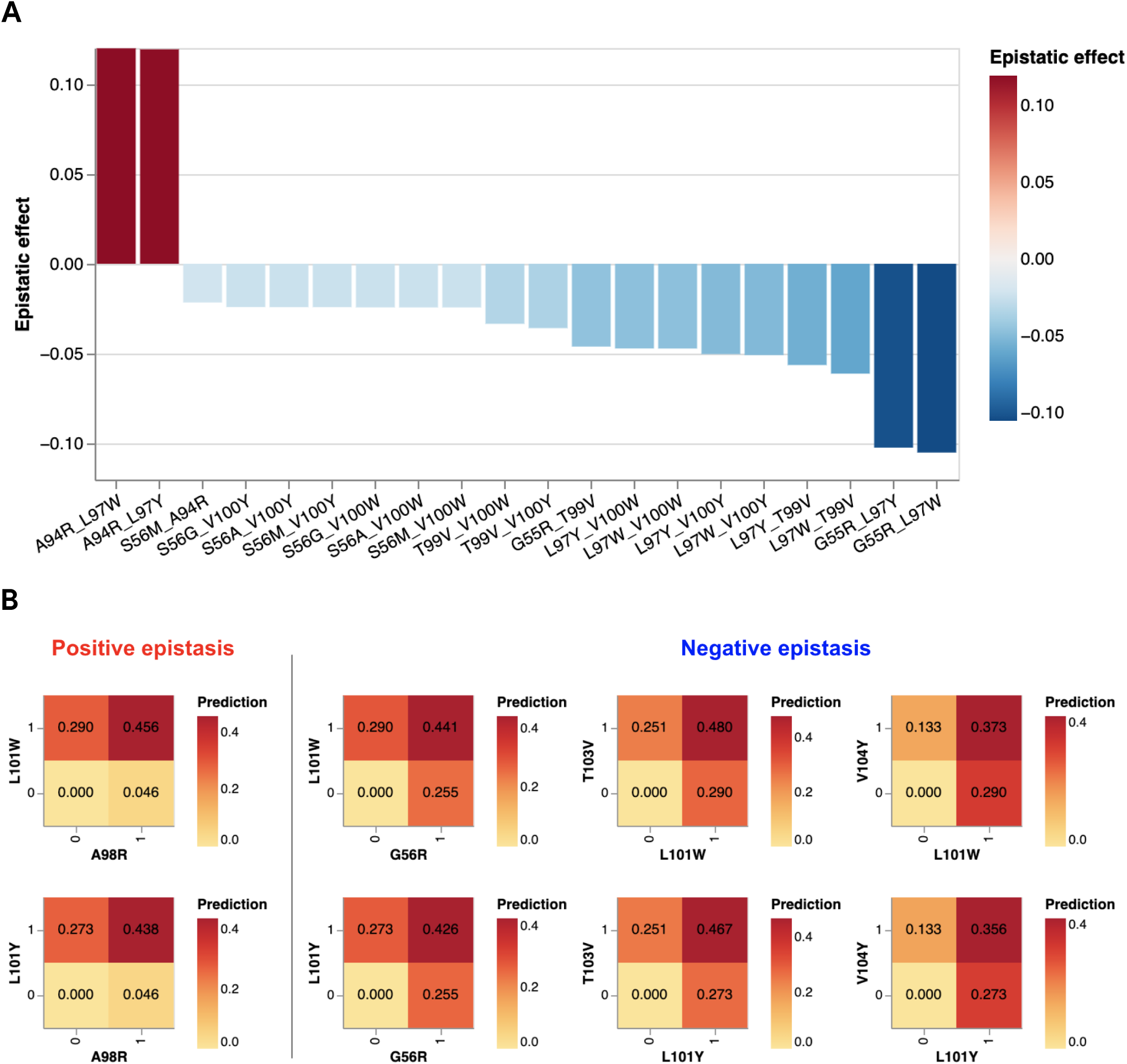
Residue pairs with the strongest predicted epistatic effect. (A) Positive and negative values denote that two mutations in combination increase or decrease binding more than in isolation, respectively. (B) Predicted single and double mutant effect sizes of residue pairs with the strongest epistatic effect. On-diagonal values denote the predicted effect of single mutants. Off-diagonal values in the top right corner corresponds to the predicted effect of double mutant. The difference corresponds to the epistatic effect size.

**Extended Data Fig. 9.**
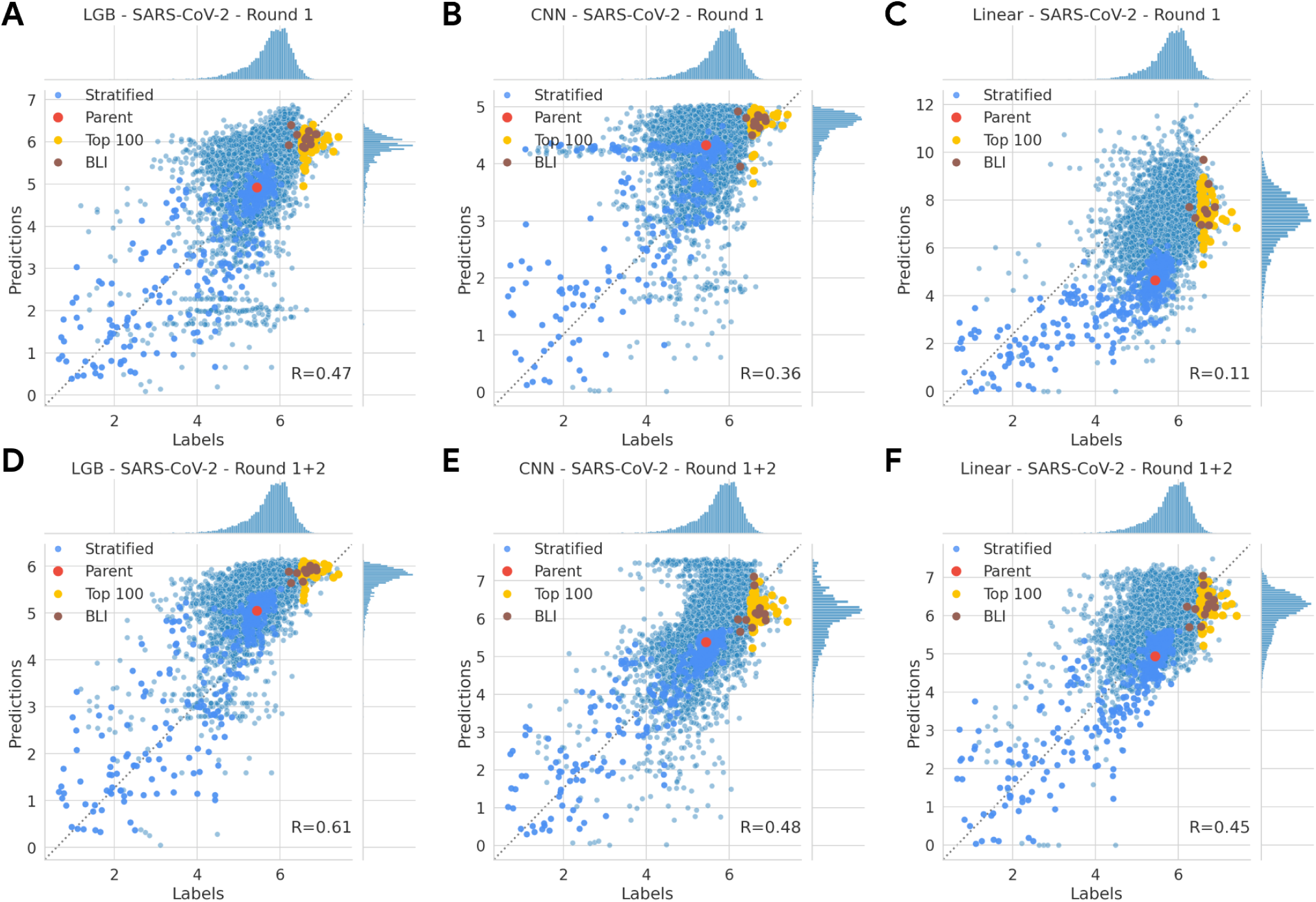
Scatterplots of model predictions vs. ground truth labels for SARS-CoV-2 and models trained on either only round 1 (top row), or round 1 and round 2 (bottom row). Each dot corresponds to one hold-sequence assayed in round 3 that was not used for model training. Labels and predictions correspond to binding affinities (the higher the better) obtained by normalizing AlphaSeq log KD values and computing the difference from the maximum observed log KD. Red: the parent sequence VHH-72; Blue: stratified sequences with both low and high AlphaSeq affinity measurements; Yellow: sequences with the highest AlphaSeq affinity measurements; Brown: sequences with the highest BLI affinity measurements. Note that panel D is identical to Figure 5A from the main text, and is reproduced here for convenience.

**Extended Data Fig. 10.**
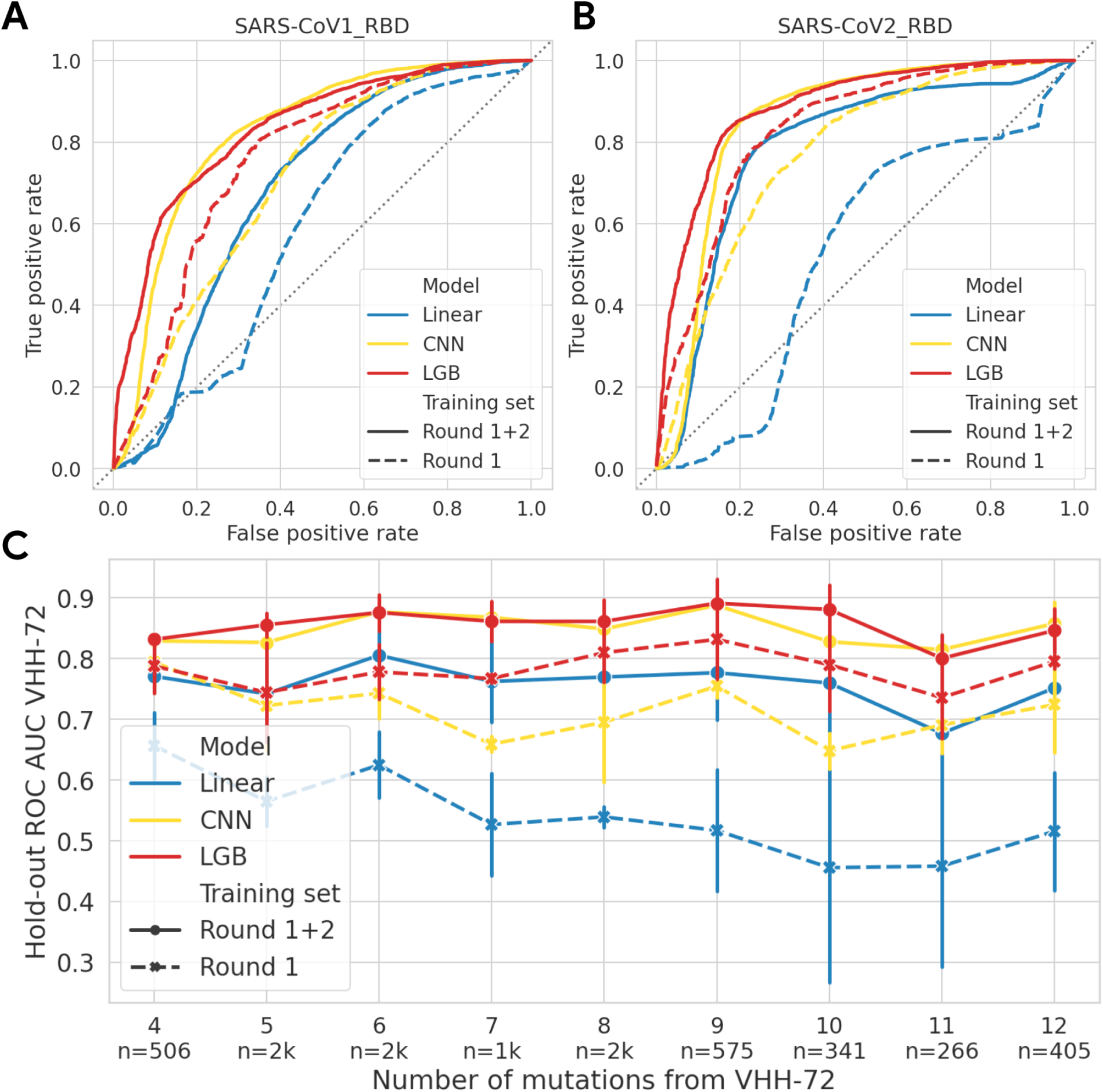
Prospective validation of model classification performance, measured using the ML-designed sequences that were experimentally tested in Round 3. (A-B) Receiver operating characteristic curves (ROC) to discriminate between sequences with a binding affinity above or below parental VHH-72 for SARS-CoV-1 and SARS-CoV-2. (C) 5-fold cross-validation ROC AUC scores stratified by the number of mutations from VHH-72.

**Extended Data Table 1.** List of CoV and CoV-related targets that were assayed by AlphaSeq in the first, second and third libraries.

| Library | Target name | is_control | control_type | Group |
| --- | --- | --- | --- | --- |
| 1 | HKU3-1 | FALSE | None | BatAsia |
| 1 | RF1 | FALSE | None | BatAsia |
| 1 | Rp3 | FALSE | None | BatAsia |
| 1 | ZC45 | FALSE | None | BatAsia |
| 1 | ZXC21 | FALSE | None | BatAsia |
| 1 | BM48-31 | FALSE | None | BatNonAsia |
| 1 | LYRa11 | FALSE | None | CoV1 |
| 1 | SARS-CoV1_RBD | FALSE | None | CoV1 |
| 1 | WIV1 | FALSE | None | CoV1 |
| 1 | Pangolin | FALSE | None | CoV2 |
| 1 | RaTG13 | FALSE | None | CoV2 |
| 1 | SARS-CoV2_RBD | FALSE | None | CoV2 |
| 1 | SARS-CoV2_RBD_G502D | FALSE | None | CoV2 |
| 1 | SARS-CoV2_RBD_N439K | FALSE | None | CoV2 |
| 1 | SARS-CoV2_RBD_N501D | FALSE | None | CoV2 |
| 1 | SARS-CoV2_RBD_N501F | FALSE | None | CoV2 |
| 1 | SARS-CoV2_RBD_R408I | FALSE | None | CoV2 |
| 1 | SARS-CoV2_RBD_S477N | FALSE | None | CoV2 |
| 1 | SARS-CoV2_RBD_V367F | FALSE | None | CoV2 |
| 1 | SARS-CoV2_RBD_V483A | FALSE | None | CoV2 |
| 1 | MERS-CoV | FALSE | None | MERS |
| 1 | MATalpha_Negative_1 | TRUE | negative | negative_control |
| 1 | MATalpha_Negative_2 | TRUE | negative | negative_control |
| 1 | MATalpha_Negative_3 | TRUE | negative | negative_control |
| 1 | MATalpha_Control_1 | TRUE | positive | positive_control |
| 1 | MATalpha_Control_2 | TRUE | positive | positive_control |
| 1 | MATalpha_Control_3 | TRUE | positive | positive_control |
| 1 | MATalpha_Control_4 | TRUE | positive | positive_control |
| 1 | MATalpha_Control_5 | TRUE | positive | positive_control |
| 1 | MATalpha_Control_6 | TRUE | positive | positive_control |
| 2 | SARS-CoV1_RBD | FALSE | None | CoV1 |
| 2 | SARS-CoV2_RBD | FALSE | None | CoV2 |
| 2 | SARS-CoV2_RBD_N501Y | FALSE | None | CoV2 |
| 2 | SARS-CoV2_RBD_N501Y+K417N+E<br>484K | FALSE | None | CoV2 |
| 2 | SARS-CoV2_RBD_N501Y+K417T+E<br>484K | FALSE | None | CoV2 |
| 2 | SARS-CoV2_RBD_R408I | FALSE | None | CoV2 |
| 2 | MATalpha_Negative_1 | TRUE | negative | negative_control |
| 2 | MATalpha_Negative_2 | TRUE | negative | negative_control |
| 2 | MATalpha_Negative_3 | TRUE | negative | negative_control |
| 2 | MATalpha_Control_1 | TRUE | positive | positive_control |
| 2 | MATalpha_Control_2 | TRUE | positive | positive_control |
| 2 | MATalpha_Control_3 | TRUE | positive | positive_control |
| 2 | MATalpha_Control_4 | TRUE | positive | positive_control |
| 2 | MATalpha_Control_5 | TRUE | positive | positive_control |
| 2 | MATalpha_Control_6 | TRUE | positive | positive_control |
| 3 | SARS-CoV1_RBD | FALSE | None | CoV1 |
| 3 | SARS-CoV2_RBD | FALSE | None | CoV2 |
| 3 | MATalpha_Negative_1 | TRUE | negative | negative_control |
| 3 | MATalpha_Negative_2 | TRUE | negative | negative_control |
| 3 | MATalpha_Negative_3 | TRUE | negative | negative_control |
| 3 | MATalpha_Control_1 | TRUE | positive | positive_control |
| 3 | MATalpha_Control_2 | TRUE | positive | positive_control |
| 3 | MATalpha_Control_3 | TRUE | positive | positive_control |
| 3 | MATalpha_Control_4 | TRUE | positive | positive_control |
| 3 | MATalpha_Control_5 | TRUE | positive | positive_control |
| 3 | MATalpha_Control_6 | TRUE | positive | positive_control |

**Extended Data Table 2.**
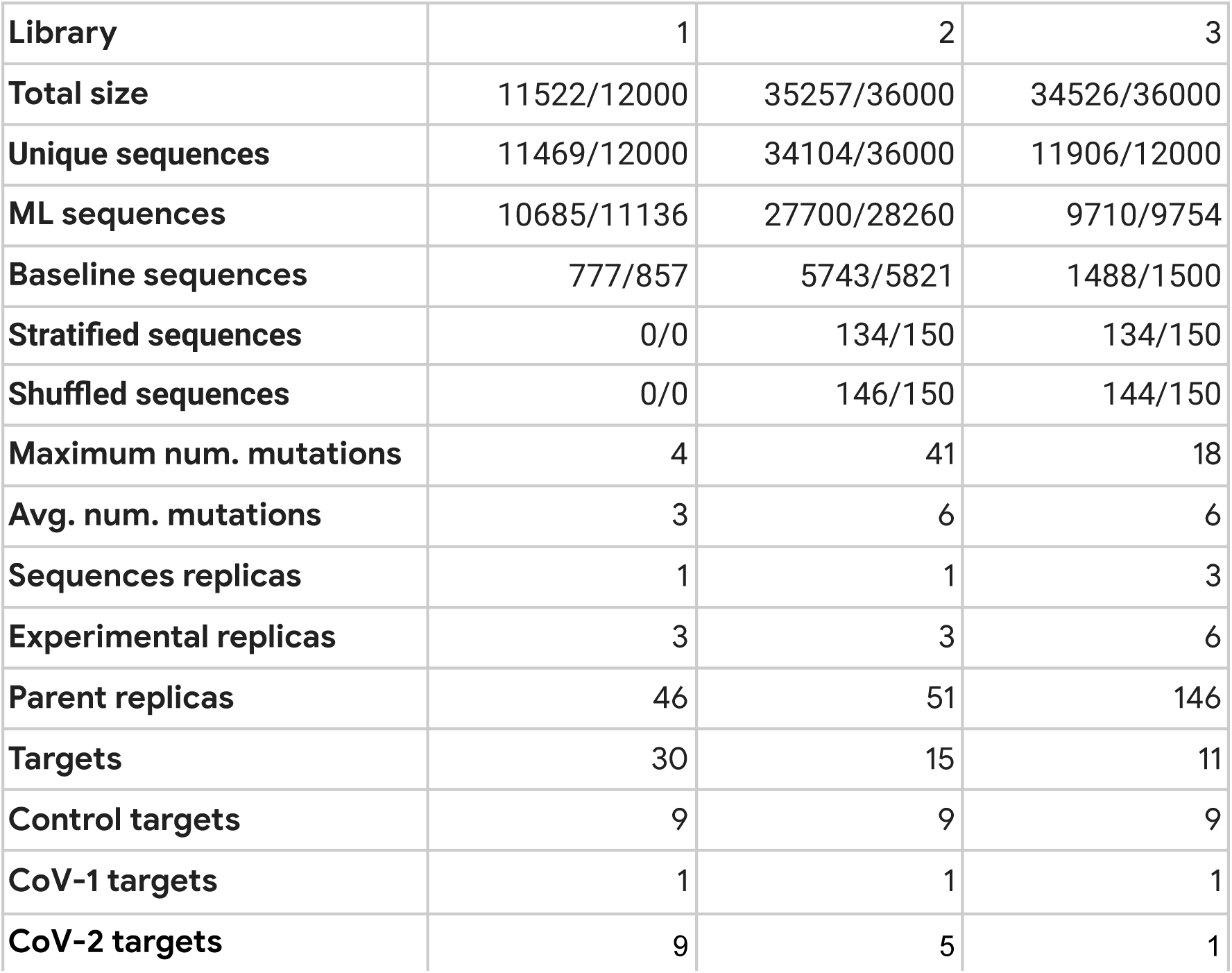
Number of sequences, sequence replicates, and SARS-CoV targets of the three designed libraries. The number of sequences with observed AlphaSeq measurements, and the total number of designed sequences is denoted before and after the forward slash, respectively. Dropouts can be the result of sequencing or experimental errors.

| Library | 1 | 2 | 3 |
| --- | --- | --- | --- |
| **Total size** | 11522/12000 | 35257/36000 | 34526/36000 |
| **Unique sequences** | 11469/12000 | 34104/36000 | 11906/12000 |
| **ML sequences** | 10685/11136 | 27700/28260 | 9710/9754 |
| **Baseline sequences** | 777/857 | 5743/5821 | 1488/1500 |
| **Stratified sequences** | 0/0 | 134/150 | 134/150 |
| **Shuffled sequences** | 0/0 | 146/150 | 144/150 |
| **Maximum num. mutations** | 4 | 41 | 18 |
| **Avg. num. mutations** | 3 | 6 | 6 |
| **Sequences replicas** | 1 | 1 | 3 |
| **Experimental replicas** | 3 | 3 | 6 |
| **Parent replicas** | 46 | 51 | 146 |
| **Targets** | 30 | 15 | 11 |
| **Control targets** | 9 | 9 | 9 |
| **CoV-1 targets** | 1 | 1 | 1 |
| **CoV-2 targets** | 9 | 5 | 1 |

